# Early and transient increase in cortical pyramidal cell excitability and delayed alteration of evoked synaptic transmission and t-SNARE proteins content in the hippocampus and neocortex of neonatal and juvenile *STXBP1* heterozygous mice

**DOI:** 10.64898/2025.11.29.691294

**Authors:** Louison Pineau, Hélène Becq, Emilie Pallesi-Pocachard, Melanie Brosset-Heckel, Najoua Biba-Maazou, Aurelie Montheil, Mathieu Milh, Pierre-Pascal Lenck-Santini, Laurent Aniksztejn

## Abstract

*De novo* mutations in the *STXBP1* gene leads to the haploinsufficiency of Munc18.1 protein in patient, and represent one of the major causes of neurodevelopmental disorders including developmental epileptic and non-epileptic encephalopathies. Given the fundamental role of this protein in vesicular exocytosis, most electrophysiological studies have focused on the impact of this haploinsufficiency on synaptic transmission, and much less on intrinsic neuronal properties. Furthermore, the possibility that the electrophysiological consequences may be dependent on the developmental stage has not yet been investigated. Here, we analyze using acute brain slices from neonatal and juvenile *STXBP1* heterozygous mice the intrinsic properties as well as spontaneous and evoked glutamatergic and GABAergic synaptic transmission in pyramidal cells located in the CA1 region of the hippocampus and in layers II/III of the motor cortex. We show that Munc18.1 deficiency has different electrophysiological consequences in neonatal (postnatal days, PND 4-7) and juvenile mice (PND30-35). The deficit of Munc18.1 leads to an increase in the intrinsic excitability of hippocampal and motor cortical pyramidal cells in neonates while in juveniles, it is evoked synaptic transmission that is affected, with a greater sensitivity of glutamatergic synapses than of GABAergic synapses in response to high-frequency electrical stimulation. However spontaneous ongoing synaptic activity mediated by glutamate and GABA receptors was unaffected at both stages of development. In addition, we performed western blot analysis and observed that Munc18.1 deficiency in STXBP1 heterozygous mice is associated with decreased t-SNARE proteins expression levels in the hippocampus and neocortex of juvenile but not neonatal mice. Therefore, Munc18.1 deficiency has multiple electrophysiological and biochemical consequences, which depend on the developmental stage. These data suggest also that an alteration in the function of some ion channels is one of the first electrophysiological consequences of the deficit of Munc18.1 and we suggest that the decrease in t-SNARE proteins expression could contribute to the normalization of pyramidal cells firing properties in juvenile.

## Introduction

Munc18.1, a protein encoded by the *STXBP1* gene, is an intrinsic component of the fusion machinery that binds to the target plasma membrane SNARE (soluble N-ethylmaleimide-sensitive factor attachment protein receptor) protein syntaxin 1, and form a trans-SNARE complex together with SNAP25 and the vesicular SNARE protein synaptobrevin that is instrumental for the docking of synaptic vesicles at the active zone, their priming and their fusion with the plasma membrane for transmitter release (Sudhöf, 2014). Deletion of the *STXBP1* gene fully abolishes synaptic transmission (Verhage et al., 2000). *De novo* mutations in the *STXBP1* gene have been identified in a variety of syndromes ranging from developmental epileptic encephalopathy with early onset (DEE) to non-epileptic encephalopathy (Saitsu et al., 2008; Di Meglio et al., 2015; Stamberger et al.,2016; Abramov et al. 2021). These mutations (missense, non-sense, frameshift, intronic, deletion) are in most of cases loss of function leading to the degradation of the protein or to its aggregation and the inability of Munc18.1 to interact with its different partners. Thus *STXBP1* related patients are haploinsufficient for functional Munc18.1 (Saitsu et al., 2012; Kovacevic et al., 2018; Van Berkel et al., 2024). The fundamental role played by Munc18.1 in transmitter exocytosis has led to the hypothesis that STXBP1-related neurodevelopmental disorders are caused by an alteration in synaptic transmission. A number of studies have been performed to investigate this issue in several preparations including neuronal cultures from a heterozygous *STXBP1* mouse model or expressing Human pathogenic variants (Toonen et al, 2006; Kovacevic et al., 2018; Orock et al., 2018; Guiberson et al., 2024); Human neurons conditionally expressing STXBP1 loss-of-function mutations (Patzke et al., 2015), Human organotypic slice culture in which STXBP1 was inactivated by RNA interference (Mc Leod et al., 2023); STXBP1 patients-IPSC derived neurons (Van Berkel et al. 2024), acute brain slices from different STXBP1 heterozygous mouse models (Miyamoto et al., 2019; Chen et al., 2020; Dos Santos et al., 2023) and in which high K^+^ or electrical stimuli, more often at high frequencies, were applied to assess vesicular exocytosis and the efficiency of synaptic transmission. While all studies in which high K^+^ was used showed impaired vesicular release under Munc18.1 deficient conditions (Orock et al., 2018; McLeod et al., 2023; Guiberson et al., 2024), opposing results were obtained regarding the response of synapses to electrical stimulation. Some studies have shown a faster and stronger rundown of the synaptic response to high frequency electrical stimulation and a weaker synaptic response to a single stimulus under Munc18.1 deficiency compared to control conditions while in other studies synaptic transmission evoked by same stimuli patterns were not affected (Toonen et al., 2006; Patzke et al., 2015; Kovacevic et al., 2018; Van Berkel et al., 2024). In acute brain slices, these investigations have mainly been carried out on synapses made between pyramidal cells of the somatosensory cortex with other pyramidal neurons or paravalbumin and somatostatin interneurons located in the same region and vice versa or with GABAergic interneurons of the striatum (Miyamoto et al.,2019; Chen et al., 2020; Dos Santos et al., 2023). These studies revealed that, even within the same structure, glutamatergic and GABAergic synapses may exhibit different sensitivity to Munc18.1 deficiency. In particular, GABAergic synapses on pyramidal cells, unlike glutamatergic synapses, respond similarly to high-frequency stimulation or sustained depolarization in wild-type and STXBP1 heterozygous mice (Chen et al., 2020; Dos Santos et al., 2023); this may have important consequences for the balance between excitation and inhibition within cortical circuits.

However, this mechanism may not be the only one involved in *STXBP1*-related diseases. In addition to synaptic transmission, Munc18.1 has been shown to indirectly affect the function of several ion channels, including voltage-gated potassium and calcium channels in heterologous cells, by preventing the interaction and inhibitory action of Syntaxin 1 on these channels (Jarvis and Zamponi, 2001; Gladisheva et al. 2004; Devaux et al., 2017). This raises the possibility that the deficiency of Munc18.1 could affect neuronal excitability. Consistent with this hypothesis, although opposing results have been obtained, an alteration of neuronal activity (an increase and a decrease) has been reported in *STXBP1* patients-IPSC derived neurons and neuronal cultures from *STXBP1* heterozygous mice, in which the discharge was assessed using fluorometric calcium imaging and multi-electrode array respectively (Van Berkel et al., 2024; Guiberson et al., 2024).

Here we performed electrophysiological recordings in acute brain slices from an *STXBP1* heterozygous mice model (Toonen et al., 2005, 2006; Kovacevic et al., 2018) and addressed the following three questions: i) Does reduced Munc18.1 expression affect the intrinsic properties of neurons? ii) Can the differential impact of Munc18.1 deficiency on glutamate and GABAergic synapses response to high frequency electrical stimulation in the somatosensory cortex can be generalized in other cortical structures? iii) Are spontaneous ongoing synaptic activities mediated by glutamate and GABA affected? To answer these questions, we recorded pyramidal cells located in the CA1 region of the hippocampus and in layers II/III of the motor cortex, two structures that are likely to participate in the generation of seizures and to the various motor and cognitive deficits that have been described in *STXBP1* heterozygous mice models (Kovacevic et al., 2018; Orock et al., 2018; Miyamoto et al., 2019; Chen et al., 2020). We conducted this study at the neonatal (Post Natal Day, PND 4-7) and, for comparison, also at juvenile (PND 30-35) stages of development. The analysis in neonates was motivated by the following reasons: i) Munc18.1 is expressed early in the developing brain (Hamada et al., 2017; Verhage et al., 2000); ii) Clinical manifestations including seizures and abnormal interictal activity in *STXBP1* patients are already observed at birth (Saitsu et al., 2008; Di Meglio et al. 2015); ii) Some electrophysiological alterations observed in other models of DEE are time limited being observed only at early stages of development but not later (Biba-Maazou et al., 2022; Reva et al., 2025; Mao et al., 2025; Wang et al., 2025); iii) The first postnatal week of life may be a critical period for the development of disease in other mouse model of DEE (Peters et al., 2005; Marguet et al., 2015). Here we show that a Munc18.1 deficit has multiple electrophysiological and biochemical consequences depending on the developmental stage.

## Methods

### Animals and genotyping

All experiments were carried out in accordance with the European Communities Council Directive of September 22, 2010 (2010/63/UE) related to laboratory animals used for research purposes. The study was approved by ethics commity of the “Ministère de l’Enseignement Supérieur de la Recherche et de l’Innovation (APAFIS#27891-2020110514571312 v2). Experiments were performed on male and female wild type and STXBP1 heterozygous C56/Bl6 mice aged 4-7 days old (neonatal mice, PND 4-7) and aged 30-35 days old (juvenile mice, PND 30-35). Mice were housed in a temperature-controlled environment with a 12 light/dark cycle and free access to food and water (INMED animal facilities).

### Slices preparation

Experiments were performed on pyramidal neurons located in deep proximal CA1 pyramidal cells of the dorsal hippocampus and in layers II/III of motor cortical slices.

Mice were decapitated after they have been euthanized by cervical dislocation. The brain was rapidly removed and placed in an oxygenated ice-cold choline solution containing (in mM): 132.5 choline chloride, 2.5 KCl, 0.7 CaCl_2_, 3 MgCl_2_, 1.2 NaH_2_PO_4_, 25 NaHCO_3_ and 8 glucose; oxygenated with 95% O_2_ and 5% of CO_2_. Coronal slices (300 μm thick) were cut using a vibratome (Leica VT1200S; Leica Microsystems, Germany) in ice-cold choline solution oxygenated with 95% O_2_ and 5% of CO_2_. Before recording, slices were incubated in an artificial cerebrospinal fluid (ACSF) solution with the following composition (in mM): 125 NaCl, 3.5 KCl, 2 CaCl_2_, 1.3 MgCl_2_, 1.25 NaH_2_PO_4_, 26 NaHCO_3_ and 10 glucose equilibrated at pH 7.3-7.4 with 95% O_2_ and 5% CO_2_ at 34 °C for 20 min and then at room temperature (22–25 °C) for at least 1 h to allow recovery. Slices were placed into the recording chamber for electrophysiological purpose where they were fully submerged and superfused with oxygenated ACSF solution at 34-35 °C at a rate of 5 ml/min.

### Electrophysiology

Pyramidal neurons were recorded under visual control with a Nikon microscope in whole cell configurations using borosilicate glass capillaries (GC 150F-15). For recordings in current-clamp mode, patch-pipettes were filled with a solution containing (in mM): 140 KMeSO_4_, 6 NaCl, 10 HEPES, 1 MgCl_2_, 4 Mg-ATP, 0.4 Na_2_-GTP. The pH was adjusted to 7.35 with KOH. The resistance of the pipettes was of 5-6 MΩ. For recordings of spontaneous synaptic activities in voltage-clamp mode, patch pipettes were filled with a solution containing (in mM): 120 Cs-gluconate, 10 CsCl, 10 HEPES, 4 Mg-ATP, 0.4 Na_2_-GTP. The pH was adjusted to pH 7.35 with CsOH. The resistance of the pipettes was of 6-7 MΩ. For recordings of evoked synaptic currents, patch pipettes were filled with a solution containing (in mM): 140 KCl, 10 HEPES, 4 Mg-ATP, 0.4 Na_2_-GTP. The pH was adjusted to pH 7.35 with KOH. The resistance of the pipettes was of 5-7 MΩ.

All reported potential values were corrected for the liquid junction potential, calculated to be ∼2 mV with the KMeSO_4_ pipette solution and ∼15 mV with the Cs-gluconate filled pipette solution.

Recordings in current-clamp mode performed to assess the intrinsic properties of neurons were carried out in the presence of 2,3-Dioxo-6-nitro-1,2,3,4-tetrahydrobenzo[*f*]quinoxaline-7-sulfonamide (NBQX, 10 µM) to block AMPA/Kainate receptors; D-2-amino-5-phosphonovalerate (D-APV, 40 µM) to block NMDA receptors; 6-Imino-3-(4-methoxyphenyl)-1(6*H*)-pyridazinebutanoic acid hydrobromide (SR 95531, Gabazine 5 µM) to block GABA_A_ receptors. All these drugs were purchased from Tocris Bioscience (Bristol, UK).

Resting membrane potential was determined in current clamp as the potential upon break-in, before any current injection. Values were corrected *post hoc* according to the formula of Tyzio et al. (2003). Action potential (AP) threshold was defined as the membrane potential at which the rate of depolarization is maximal. All measurements were filtered at 3 KHz using an EPC10 amplifier (HEKA Electronik, Germany) and sampled at 10 kHz. Data were analyzed off-line using Clampfit (Molecular Devices), miniAnalysis (Synaptosoft), Origin 9 (Origin Lab) and Prism 6 software (GraphPad).

Only cells with a stable resting membrane potential more negative than –55 mV were used in this study. All measurements in current-clamp mode were performed from a membrane potential of – 70 mV and if necessary, current was injected during the experiment to keep this value constant.

To calculate the membrane input resistance (Rm), voltage responses to the injection of five depolarizing and hyperpolarizing current steps, with an increment of +/−5 pA and applied during 500 msec, were fit with a linear function to yield the slope value.

To calculate the membrane time constant (τ_m_), the voltage response to the injection of a hyperpolarizing current step of –20 pA for 500 msec was fitted with a single exponential function (Origin 9) to yield the tau value. Membrane capacitance (Cm) was then calculated according to the equation C_m_ = τ_m_/R_m_.

Action potentials were elicited by injection of short 10 msec current steps command of 50 pA increment to assess their properties, and to assess neuronal excitability, by injection of long 1 sec depolarizing current steps of 10 to 150 pA (in 10 pA increments) in pyramidal cells from animal aged one week and of 20-300 pA (in 20 pA increments) from older animals.

### Synaptic transmission evoked by electrical stimulation

GABAR and AMPAR-mediated postsynaptic current (PSC) evoked by electrical stimulation were both recorded with a KCl filled pipette solution (see above). GABAR-PSCs were recorded in continuous presence of CNQX (10 µM) and D-APV (40 µM). AMPAR-mediated PSC were recorded in continuous presence of gabazine (10µM) and D-APV (40 µM) and with external solution containing high Mg^2+^ (6mM), High Ca^2+^ (4mM) to prevent polysynaptic events that would be favor with the use of gabazine. Both GABAR and AMPAR-PSCs were recorded at –70 mV. The protocol consisted of inducing five to ten PSC by electrical stimulation at 0.1 Hz, followed by stimulation at 10 Hz or 30 Hz for 5 seconds and then again by stimulations at 0.1 Hz. This procedure was repeated one to three times in each cell and the responses were averaged. The stimulation intensity was adjusted to obtain an amplitude response ranging between 50 and 200 pA. The amplitude of the evoked PSC was expressed as a percentage of the PSC evoked at 0.1 Hz before the 10 or 30 Hz stimulation (average of the 5 PSCs preceding the stimulation train plus the first PSC of the train).

The size of the readily releasing pool of vesicles (RRP) was estimated in each experiment from the graph representing the cumulative PSC (in pA) versus the stimulus number and by fitting a straight line to stimuli 30-50 (at 10 Hz) or 100-150 (at 30 Hz) and then back-extrapolating this line to the y-axis. The y-intercept indicates the total current that can be generated after all vesicles in the pool are depleted before being replenished (Schneggenburger et al., 1999). The probability of vesicular release was calculated by dividing the value of the first PSC of the train by the estimated value of the size of RRP.

### Spontaneous synaptic activity

Spontaneous synaptic activities were recorded with a CsGlu filled pipette solution (see above). sGABA-PSCs were recorded at 0 mV (LJP corrected), the reversal potential GluR-PSC were recorded at –60 mV, a value close to the estimated reversal potential of GABAR-PSC (–70mV). Synaptic events were recorded during 3-5 min at the two membrane potentials.

### Western blot

Mice aged 4 days and 30 days old were decapitated after they have been euthanized by cervical dislocation, the brains were quickly removed and placed in the choline chloride solution at 4°C. The hippocampus and cortex were dissected and homogenized by mechanical dissociation (26G syringe) in RIPA buffer (Thermo Fisher Scientific) supplemented with a protease and phosphatase inhibitor cocktail (Thermo Fisher Scientific), according to the manufacturer’s instructions. Protein concentrations were determined using the BCA Protein Assay Kit (Thermo Fisher Scientific). Equal amounts of protein (15 µg for P4 and 26 µg for P30) were denatured at 95 °C for 5 min in Laemmli buffer, separated by SDS–PAGE (bis-tris 4-12%, Thermo Fisher Scientific), and transferred onto nitrocellulose membranes using a wet electrotransfer system. Membranes were blocked for 1 h at room temperature in TBS-T (Tris-buffered saline, 0.1 % Tween-20) containing 3 % bovine serum albumin (BSA), and then incubated overnight at 4 °C with the following primary antibodies diluted in blocking buffer: anti-Munc18 (GTX114899, GeneTex, 1:2000), anti-Syntaxin1A (AF7237, R&D, 1:1000), anti-Syntaxin1B (110011, Synaptic Systems, 1:2000), anti-SNAP25 (111002, Synaptic Systems, 1:5000), and anti-Vinculin (14-9777-82, Invitrogen, 1:1000). After three washes in TBS-T, membranes were incubated for 1 h at room temperature with HRP-conjugated secondary antibodies (anti-rabbit, anti-goat, or anti-mouse IgG, 1:10000; Thermo Fisher Scientific). Protein bands were visualized using enhanced chemiluminescence (ECL; WBKLS0500, Millipore) and imaged with a G-Box (Syngene). Densitometric analyses were performed using ImageJ software, and protein levels were normalized to Vinculin as a loading control.

### Statistical methods

Data shown in graphs are represented as mean ± SEM. The Student t-test was used to compare means of two groups when the data distribution was normal. When the normality test failed or the number of experiments was < 15, we used the non-parametric Mann – Whitney test for two independent samples. The normality was assessed using Shapiro-Wilk test. Those statistical analyses were performed using GraphPad Prism v9 software. To statistically evaluate the difference between two groups of curves, we applied a non-parametric two-way ANOVA with repeated measures (two-way RM-ANOVA), in which the response (Y axis) from the same cell across the discrete values of the X axis were taken as repeated measurements. Statistical significance was indicated by asterisks: * when p < 0.05; ** when p < 0.01 and *** when p < 0.001. For better readability, number of cells, animal used and statistics (only when there is a significant difference) are indicated in the legends.

## Results

### Time limited increase of cortical pyramidal cells excitability in STXBP1 heterozygous mice

The first question we wanted to answer was whether a reduced expression of Munc18.1 affected the intrinsic properties of neurons.

We recorded CA1 pyramidal cells and limited our analysis to cells located in the deep (*closer to stratum oriens*) proximal (closer to CA2) layer of the dorsal hippocampus in order to limit as much as possible the heterogeneity of electrophysiological properties of pyramidal cells that exist along the dorsal-ventral, proximal-distal and deep-superficial axes of CA1 (Cembrowski and Spruston, 2019).

At PND 4-7, the deficit of Munc18.1 did not affect significantly cell resting membrane potential (Vm); input resistance (Rm), membrane time constant (τm) and cells capacitance (Cm) (Fig. 1Ae). We then analyzed the properties of a single action potential (AP) elicited by a short (10 msec) depolarizing current steps command of 50 pA increment. We observed that the AP amplitude was slightly but significantly increased (on average by ∼4 mV) in pyramidal cells of *STXBP1* heterozygous mice (*STXBP1^+/−^*mice) compared to pyramidal cells of wild-type mice (WT mice) whereas the rheobase, the AP threshold, and AP halfwidth were not affected (Fig. 1Aa,d). We also quantified the number of APs elicited by depolarizing current steps command of 10 pA increment from 10 to 150 pA during 1 s and measured the frequency of the discharge during the first 200 msec of the steps (initial frequency, iF) and during the last 200 msec of the steps (final frequency, fF) (see also Biba-Maazou et al., 2022). Compared to WT cells, the deficit of Munc18.1 led to an increase in the number of AP elicited by current steps with intensity > 40 pA associated with an increase of the fF but not iF /current relation (Fig.1Ab,c).

At PND 30-35 the properties of single AP and neuronal discharge were not significantly different in WT and *STXBP1^+/−^* pyramidal cells (Fig. 1Ba,b). In fact this lack of effects of Munc18.1 deficiency was observed already at PND19-21 (n = 26 cells/6 WT and n = 25 cells/6 STXBP1^+/−^postweaning mice, data not shown).

Similar electrophysiological consequences were observed in pyramidal cells of layers II/III of the motor cortex. Thus at PND 4-7, we found that the amplitude of a single action potential elicited by short depolarizing current steps command was also slightly but significantly increased in STXBP1^+/−^ cells compared to WT cells (on average by ∼5mV; Fig.2Aa,d). In addition, we observed in STXBP1^+/−^ pyramidal cells a significant increase in the number of AP elicited by long depolarizing current steps command which was associated with an increase of the final but not the initial frequency and a large leftward shift of the fF/I relation (Fig. 2Ab,c). These effects were neither observed at P30-35 (Fig.2B) nor at P19-21 (n =17 cells/3 WT and n = 15 cells/4 STXBP1^+/−^postweaning mice, data not shown).

**Figure 1:**
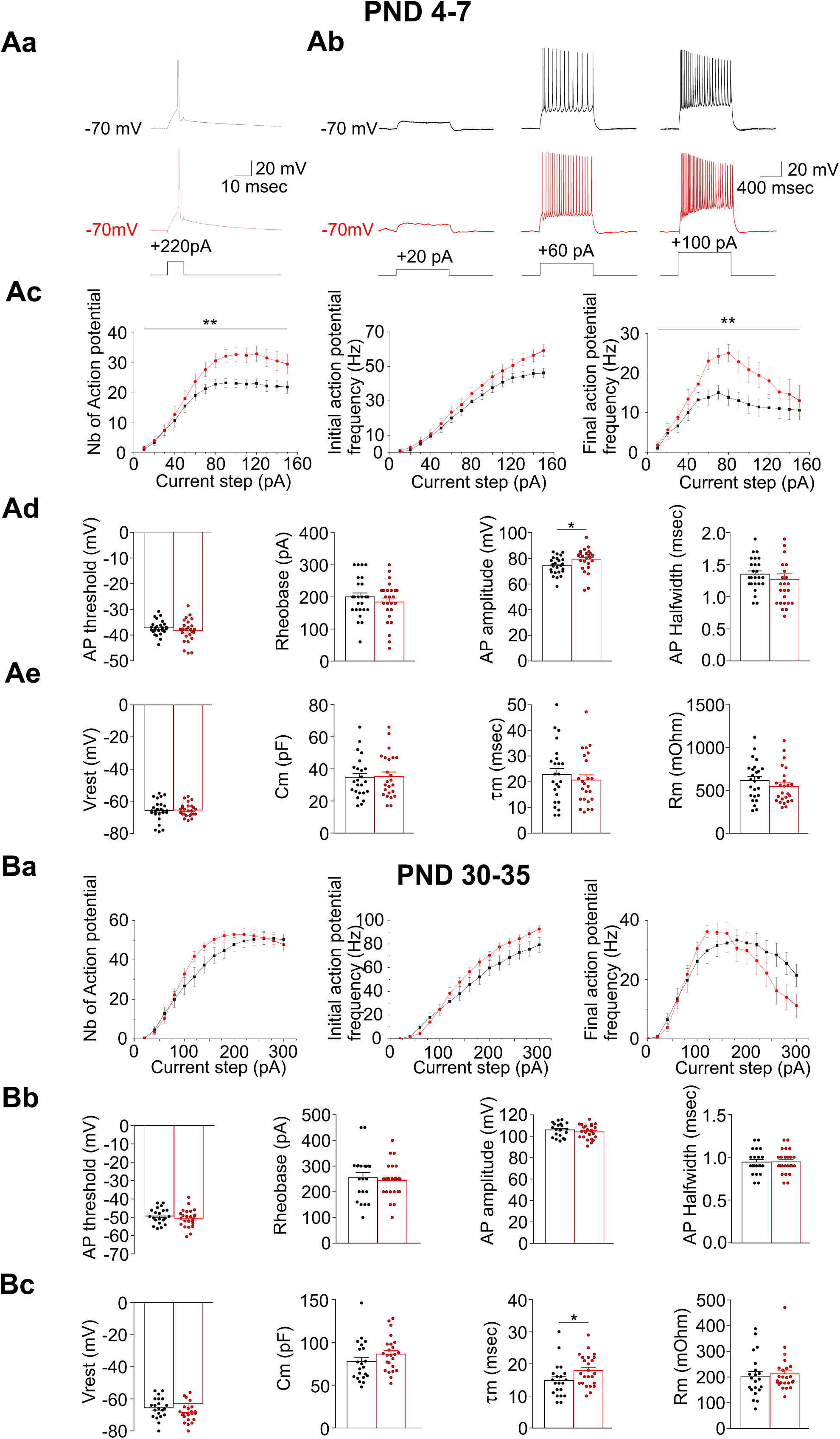
Consequences of Munc18.1 deficiency on neuronal firing and intrinsic properties of CA1 pyramidal cells of the deep proximal layer of the dorsal hippocampus recorded at PND 4-7 and PND 30-35. **Aa**) Action potential (AP) evoked by a depolarizing current step of 220 pA during 10 msec recorded in pyramidal cells from a WT mouse (black trace) and a *STXBP1^+/−^*mouse (red trace) aged 7 days old. **Ab)** Representative voltage responses to the injection of three depolarizing current steps of 20, 60 and 100 pA applied during 1sec in pyramidal cells from a WT mouse (top traces, black) and from a *STXBP1^+/−^* mouse (bottom red traces) aged 6 days old. **Ac**) Graphs quantifying (mean ± SEM), at PND 4-7, the number of action potentials elicited by the injection of depolarizing current steps (left graph), the initial frequency (measured during the first 200 msec of the steps, middle graph) and the final frequency (measured during the last 200 msec of the steps, right graph) of the discharge. Black: n = 25 cells /7 WT mice; Red: n = 25 cells/ 6 STXBP1^+/−^ mice. Statistics: AP/pA relation ** p = 0.0082, two-way repeated-measures Anova calculated from the starting point of the curve to the end; final AP frequency/pA relation: **p = 0.01 two-way repeated-measures Anova calculated from the starting point of the curve to the end. **Ad**) Boxplots quantifying the action potential (AP) threshold, the rheobase, the AP amplitude, and AP halfwidth and elicited by depolarizing current steps applied during 10 msec in WT and *STXBP1^+/−^* cells. Black: n = 25 cells /7 WT mice; Red: n = 25 cells/ 6 *STXBP1^+/−^* mice. Statistics for AP amplitude: *p = 0.015; Mann and Whitney test, Mean ± SEM in WT cells: 74.2 ± 1.4 mV and 78.8 ± 1.8 mV. **Ae**) Boxplots quantifying the resting membrane potential (Vm), the capacitance (Cm), the membrane time constant (τm) and input resistance (Rm) of WT cells and *STXBP1^+/−^* cells. **Ba-c)** Same as Ac-e but in cells recorded in mice aged between P30-35. Black: n = 21 cells/ 7 WT mice; Red: 25 cells/ 7 *STXBP1^+/−^* mice.

**Figure 2:**
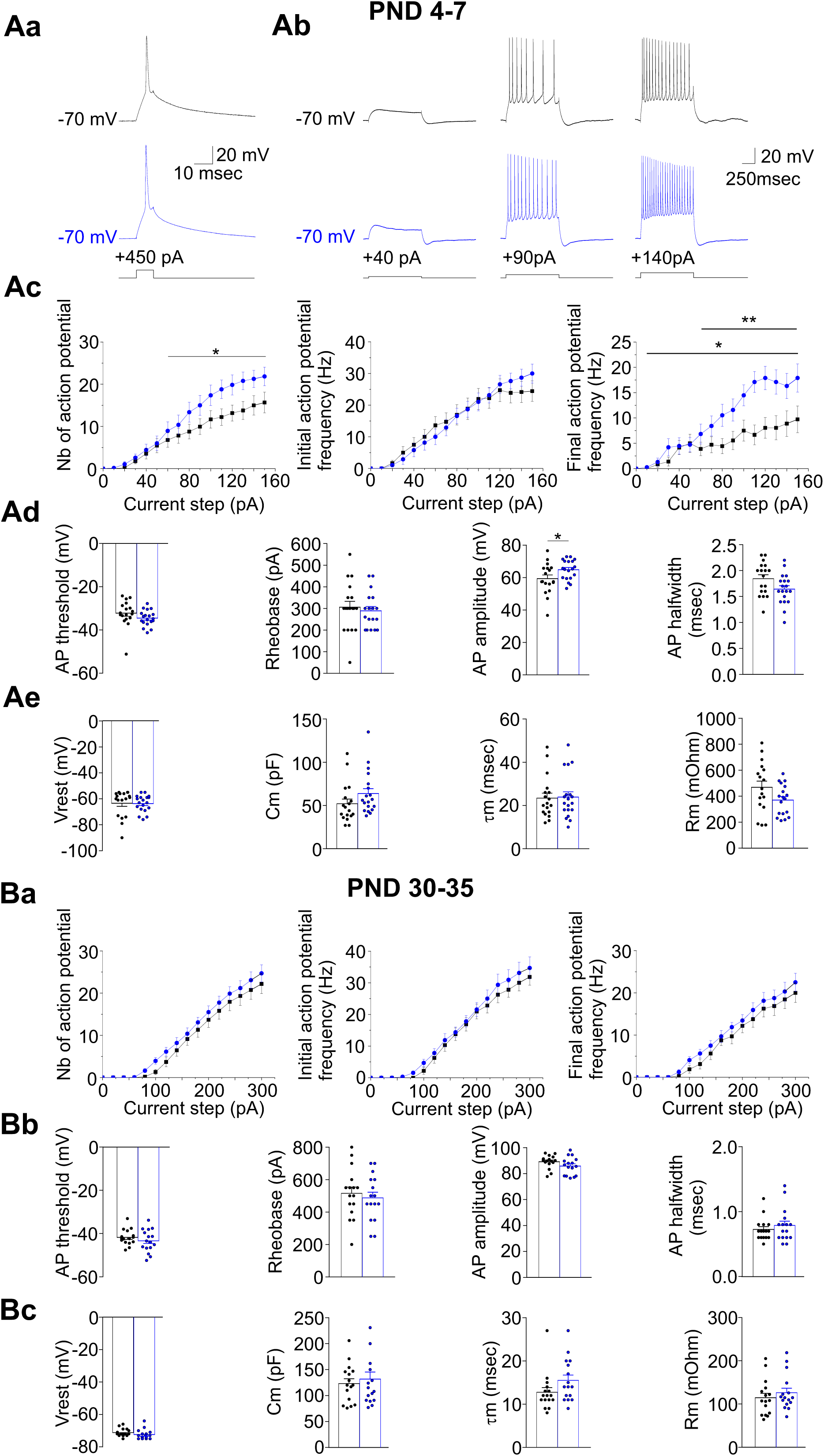
Consequences of Munc18.1 deficiency on neuronal firing and intrinsic properties of pyramidal cells located in the layers II/III of the motor cortex and recorded at PND 4-7 and PND 30-35. **Aa**) Action potential (AP) evoked by a depolarizing current step of 450 pA during 10 msec recorded in pyramidal cells from a WT mouse (black trace) and a *STXBP1^+/−^* mouse (blue trace) aged 6 days old. **Ab)** Representative voltage responses to the injection of three depolarizing current steps of 40, 90 and 140 pA applied during 1sec in pyramidal cells from a WT mouse (top traces, black) and from a *STXBP1^+/−^* mouse (bottom traces, blue) aged 6 days old. **Ac**) Graphs quantifying (mean ± SEM), at PND 4-7, the number of action potentials elicited by the injection of depolarizing current steps (left graph), the initial frequency (measured during the first 200 msec of the steps, middle graph) and the final frequency (measured during the last 200 msec of the steps, right graph) of the discharge. Black: n = 18 cells /5 WT mice; Blue: n = 19 cells/ 4 *STXBP1^+/−^*mice. Statistics: AP/pA relation * p = 0.049 two-way repeated-measures Anova calculated from currents steps of 60 pA to the end; final AP frequency/pA relation: *p = 0.012 two-way repeated-measures Anova calculated from the starting point of the curve to the end; **p = 0.0075 calculated from +60 pA to the end. **Ad**) Boxplots quantifying the action potential (AP) threshold, the rheobase, the AP amplitude, and AP halfwidth and elicited by depolarizing current steps applied during 10 msec in WT and *STXBP1^+/−^* cells. Black: n = 18 cells /5 WT mice; Blue: n = 19 cells/ 4 *STXBP1^+/−^* mice. Statistics for AP amplitude: *p = 0.043; Student t-test. Mean ± SEM = 59.4 ± 2.2 mV in WT cells and 64.8 ± 1.4 mV in *STXBP1^+/−^*cells. **Ae**) Boxplots quantifying the resting membrane potential (Vm), the capacitance (Cm), the membrane time constant (τm) and input resistance (Rm) and of wild type cells and *STXBP1^+/−^* cells. **Ba-c)** Same as Ac-e but in cells recorded in mice aged between P30-35. Black: n = 16 cells/ 4 WT mice; blue 16 cells/3 *STXBP1^+/−^* mice.

Therefore, Munc18.1 deficiency leads to increased intrinsic excitability of cortical pyramidal cells, but this electrophysiological alteration is transient and only observed during the neonatal period.

Delayed and differential sensitivity to Munc18.1 deficiency of GABAergic and glutamatergic synapses during high-frequency electrical stimulation

Next, we investigated the impact of Munc18.1 deficiency on gabaergic and glutamatergic synaptic transmission in CA1 and motor cortical layers II/III pyramidal cells in response to high frequency electrical stimulation.

In CA1, AMPA and GABA receptors-mediated postsynaptic currents (AMPAR-PSC and GABAR-PSC) were evoked using a bipolar stimulating electrode positioned in the stratum radiatum in order to stimulate the Schaffer collaterals and GABAergic interneurons located in this layer. As illustrated in Figure 3Aa in WT cells at PND4-7, the stimulation at 10 Hz during 5 sec produced a slight initial increase in the amplitude of AMPAR-PSC, which gradually decreased during the train, reaching values at the end of the train close to those recorded during the 0.1Hz pre-train period. Importantly, at this stage no significant difference in PSC amplitude was found at any time during the 10 Hz stimulation in *STXBP1^+/−^* cells compared to WT cells (Fig. 3Aa). After an initial increase in the amplitude of the AMPAR-PSC, the 30 Hz stimulation induced a progressive decrease of approximately 50% of the response at the end of the train, which is reversible after the restoration of the 0.1 Hz stimulation. As with the stimulation at 10 Hz, there was no significant difference between WT and *STXBP1^+/−^* cells in the temporal evolution and in the magnitude of AMPAR-PSC depression during the stimulation at 30Hz (Fig.3Ab).

We then performed same experiments in juvenile mice. A different situation prevailed during this period compared to neonatal period. Firstly, we observed in WT cells that the AMPAR-PSCs evoked at 10 Hz were amplified throughout the duration of the train of stimulation, and that at 30 Hz, after an initial enhancement, the responses decreased, reaching values at the end of the train that were close to the control values (Fig. 3B). Secondly, compared to WT cells, we observed in STXBP1^+/−^ cells: i) at 10 Hz, a larger and more significant increase in the amplitude of AMPAR-PSC of about 30% during the first 3 seconds of the train, followed by a rapid decrease of the responses reaching values similar to those of WT cells (Fig.3Ba); ii) at 30 Hz, a larger increase in AMPA-PSCs during the first second of the train (on average by about 30%), followed by a faster decrease of the responses which, finally, at the end of the train, was depressed (Fig.3Bb).

**Figure 3:**
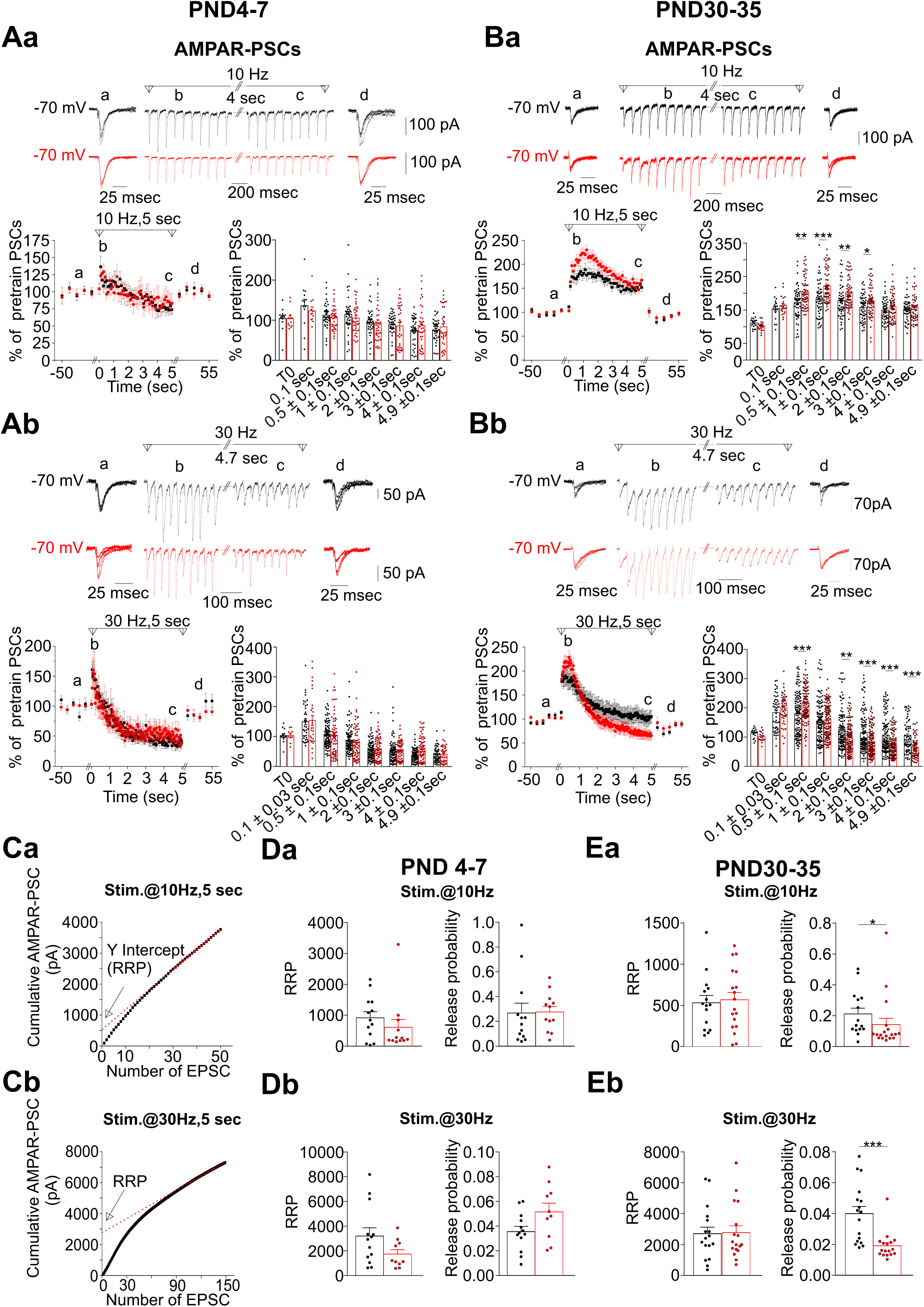
Consequences of Munc18.1 deficiency on AMPA receptors mediated postsynaptic currents (AMPAR-PSCs) in CA1 pyramidal cells and evoked by electrical stimulation applied at 10 and 30 Hz in the *stratum radiatum*. **Aa**) AMPAR mediated postsynaptic currents (AMPAR-PSCs) recorded in CA1 pyramidal cells from WT (black) and *STXBP1^+/−^* mice (red) aged 7 days old. Top: (a) 5 superimposed PSCs evoked at 0.1Hz before the stimulation at 10Hz; (b,c) PSCs during the first and the last second of the stimulation at 10 Hz stimulation (applied during 5 sec); (d) 5 superimposed PSCs evoked at 0.1Hz after the stimulation at 10Hz. Below on the left: Graph quantifying, at PND 4-7, the amplitude of PSCs (mean ± SEM), expressed as the percentage of 5 PSCs evoked at 0.1Hz before the stimulation at 10 Hz plus the first PSC of the train. Black: n = 13 cells / 3WT mice in which the stimulation protocol was applied 33 times; Red: n = 12 cells, /4 *STXBP1^+/−^* mice in which the stimulation protocol was applied 33 times. On the right: Boxplots showing average values in each cells of relative AMPAR-PSCs during the stimulation at 10 Hz and grouped at times indicated in abscissa. **Ab)** Same as Aa but for stimulation applied at 30Hz during 5 sec. Black: n = 13 cells / 3WT mice in which the stimulation protocol was applied 31 times; Red: n = 10 cells, /4 *STXBP1^+/−^* mice in which the stimulation protocol was applied 30 times. **Ba**) Same as Aa but for AMPAR-PSCs evoked by stimulation at 10 Hz in pyramidal cells from juvenile mice aged 30-35 days old. Traces of from mice aged 30 days old. Black: n = 18 cells/4 WT mice in which the stimulation protocol was applied 52 times; Red: n = 20 cells/4 *STXBP1^+/−^* mice in which the protocol was applied 55 times. Statistics at t = 0.5 ± 0.1 sec: ** p = 0.0013 Mann-Whitney test, n = 54 WT AMPAR-PSC, mean ± SEM = 171.5 ± 6.4% (expressed in % of PSCs recorded before the stimulation at 10Hz) and n = 60 *STXBP1^+/−^* AMPAR-PSCs mean = 203.8 ± 5.8 %. Statistics at t = 1 ± 0.1 sec: ***p < 0.001 Mann-Whitney test, n = 54 WT AMPAR-PSCs, mean ± SEM = 182.6 ± 6.9% and n = 60 *STXBP1^+/−^* AMPAR-PSCs, mean ± SEM= 222 ± 6.2 %. Statistics at t = 2 ± 0.1 sec: ** p = 0.0065 Mann-Whitney test, n = 54 WT AMPAR-PSCs, mean ± SEM = 178.3 ± 6.3 % and n = 60 *STXBP1^+/−^* AMPAR-PSCs, mean ± SEM = 201.6 ± 5.6%. Statistics at t = 3 ± 0.1 sec: * p= 0.011, Student t –test n = 54 WT AMPAR-PSCs, mean ± SEM = 157.4 ± 5.1% and n = 60 *STXBP1^+/−^* AMPAR-PSCs, mean ± SEM = 177.6 ± 5.9 %. **Bb)** Same as Ba but for stimulations applied at 30 Hz during 5 sec. Black: n = 17 cells/ 4 WT mice in which the stimulation protocol was applied 45 times; Red: n = 17 cells / 4 *STXBP1^+/−^* mice in which the stimulation protocol was applied 47 times. Statistics at t = 0.5 ± 0.1 sec: ***p = 0.0002 Mann-Whitney test, n = 119 WT AMPAR-PSCs, mean ± SEM = 180.6 ± 6.5% and n = 119 *STXBP1^+/−^* cells, mean ± SEM = 214.9 ± 5.2 %. Statistics at t = 2 ± 0.1 sec: ** p = 0.0056 Mann-Whitney test, n = 119 WT AMPAR-PSCs, mean ± SEM = 125.8 ± 6.2% and n = 119 *STXBP1^+/−^*AMPAR-PSCs, mean ± SEM = 99.9 ± 4.5%. Statistics at t= 3 ± 0.1 sec: ***p = 0.0001 Mann-Whitney test, n = 119 WT AMPAR-PSCs, mean ± SEM = 115.3 ± 6.1 % and n = 119 *STXBP1^+/−^* AMPAR-PSCs, mean ± SEM = 83.3 ± 0.9 %. Statistics at t = 4 ± 0.1 sec: ***p < 0.0001 Mann-Whitney test, n = 119 WT AMPAR-PSCs, mean ± SEM = 107.3 ± 5.6 % and n = 119 *STXBP1^+/−^* AMPAR-PSCs, mean ± SEM = 72.8 ± 3.6 %. Statistics at t = 4.9 ± 0.1 sec: ***p < 0.0001 Mann-Whitney test, n = 68 WT AMPAR-PSCs, mean ± SEM = 104.2 ± 6.4% and n= 66 *STXBP1^+/−^* AMPAR-PSCs, mean ± SEM = 66.2 ± 4.2 %.**C)** Example of cumulative curves of AMPAR-PSCs amplitude at 10 Hz (**a**) and 30Hz (**b**). The size of the readily releasing pool of synaptic vesicles (RRP) was estimated after fitting a straight line to stimuli 30-50 (at 10 Hz) or 100-150 (at 30 Hz) and then back-extrapolating this line to the y-axis (red dash line and arrow). **Da)** RRP in Schaffer collaterals of hippocampal slices from neonatal WT and *STXBP1^+/−^* mice and probability of glutamate release estimated at 10Hz (Black: n= 13 cells/ 3 WT mice; red: n = 12 cells/4 *STXBP1^+/−^* mice). The probability of transmitter release was calculated by dividing the amplitude of the first EPSC of the train by the RRP value. **Db)** RRP and probability of glutamate release estimated at 30 Hz (Black: n = 13 cells / 3WT mice; red: n = 10 cells/4 *STXBP1^+/−^* mice). **Ea)** RRP and probability of glutamate release estimated at 10 Hz in Schaffer collaterals of hippocampal slices from juvenile mice (Black: 15 cells/4 WT mice and n = 18 cells/ 4 *STXBP1^+/−^* mice). Statistics for the probability of glutamate release: * p = 0.011, Mann and Whitney test, mean ± SEM = 0.21 ± 0.04 in WT and 0.14 ± 0.04 in *STXBP1^+/−^*Schaffer collaterals. **Eb**) RRP and probability of glutamate release estimated at 30 Hz (Black: n = 17 cells/4 WT mice; red: n = 17 cells/4 *STXBP1^+/−^* mice). Statistics probability of glutamate release: * p = 0.0001, Mann and Whitney test, mean ± SEM = 0.04 ± 0.005 in WT and 0.02 ± 0.002 in STXBP1^+/−^ Schaffer collaterals.

We also estimated the size of the readily releasable pool of vesicles (RRP, docked and primed vesicles in the active zone that are ready for fusion and exocytosis after an action potential) and of the release probability (Pr) at Schaffer collateral synapses from the synaptic responses evoked at 10 and 30Hz in neonatal and juvenile developmental stages (see Material and Methods and Fig.3C). We found that the size of RRP did not differ in WT and *STXBP1^+/−^* neonatal or juvenile mice (Fig. 3Da, Ea). In contrast, we observed a significant decrease in the Pr of glutamate release in juvenile but not neonatal *STXBP1^+/−^* mice compared to WT mice (Fig. 3Db, Eb).

We performed the same analysis for GABAR-PSC in other series of experiments. Stimulation at 10 Hz generated a rapid and reversible depression of GABAR-PSCs in WT cells of approximately 45% at the end of the train in both neonatal and juvenile WT mice (Fig.4Aa, Ba). GABAR-PSCs were similarly affected in *STXBP1^+/−^* cells by the 10Hz stimulation. As expected, the stimulation at 30Hz produced a faster and more important depression of the synaptic responses in both WT and *STXBP1^+/−^*cells than at 10 Hz particularly at PND 4-7 (Fig. 4Ab). The level of depression reached at the end of the train was however very slightly but significantly less important (by about 7 %) in the neonatal STXBP1^+/−^ cells than in the WT cells but not different in juvenile mice (Fig. 4Ab, Bb). The RRP and Pr values estimated at 10 Hz and 30 Hz were not different in CA1 GABAergic terminals of neonatal and juvenile WT and heterozygous mice (Fig. 4Ac, Bc).

In the motor cortex, AMPAR– and GABAR-PSCs were evoked by a stimulating electrode positioned in the layers II/III. Stimulations at 10 Hz and 30 Hz produced a reversible depression of AMPAR-PSCs (Fig.5) and of GABAR-PSCs (Fig.6) in WT pyramidal cells from neonatal and juvenile mice. At PND4-7, the time course and the magnitude of the depression of both AMPAR– and GABAR-PSCs were not significantly different in *STXBP1^+/−^* cells compared to WT cells (Fig.5A,6A). At PND 30-35 the magnitude of AMPAR-PSCs depression was larger in *STXBP1^+/−^* cells by ∼15 to 10% (from the beginning to the end of the train respectively) while at 30 Hz the depression was more pronounced only during the first second of the train (by ∼ 20% on average) (Fig.5Ba,b). GABAR-PSCs in *STXBP1^+/−^*pyramidal cells also undergo significant larger depression when stimulated at 10Hz and 30Hz than in WT cells although the difference in depression level between the 2 groups of cells was small (∼5 to 10% at 10Hz from the beginning to the end of the train and ∼5% at 30Hz) (Fig.6Ba,b). As in the hippocampus, the estimated size of RRP of the stimulated glutamatergic synaptic terminals of the layers II/III was similar in WT and *STXBP1^+/−^*mice (Fig.5Ac, Bc). There was in contrast a significant increase in the estimated Pr of glutamate in juvenile but not in neonatal *STXBP1^+/−^*mice compared to WT mice (Fig.5Ac,Bc). RRP and Pr of the stimulated GABAergic synaptic terminals were unaffected by the deficit of Munc18.1 (Fig. 6Ac,Bc).

**Figure 4:**
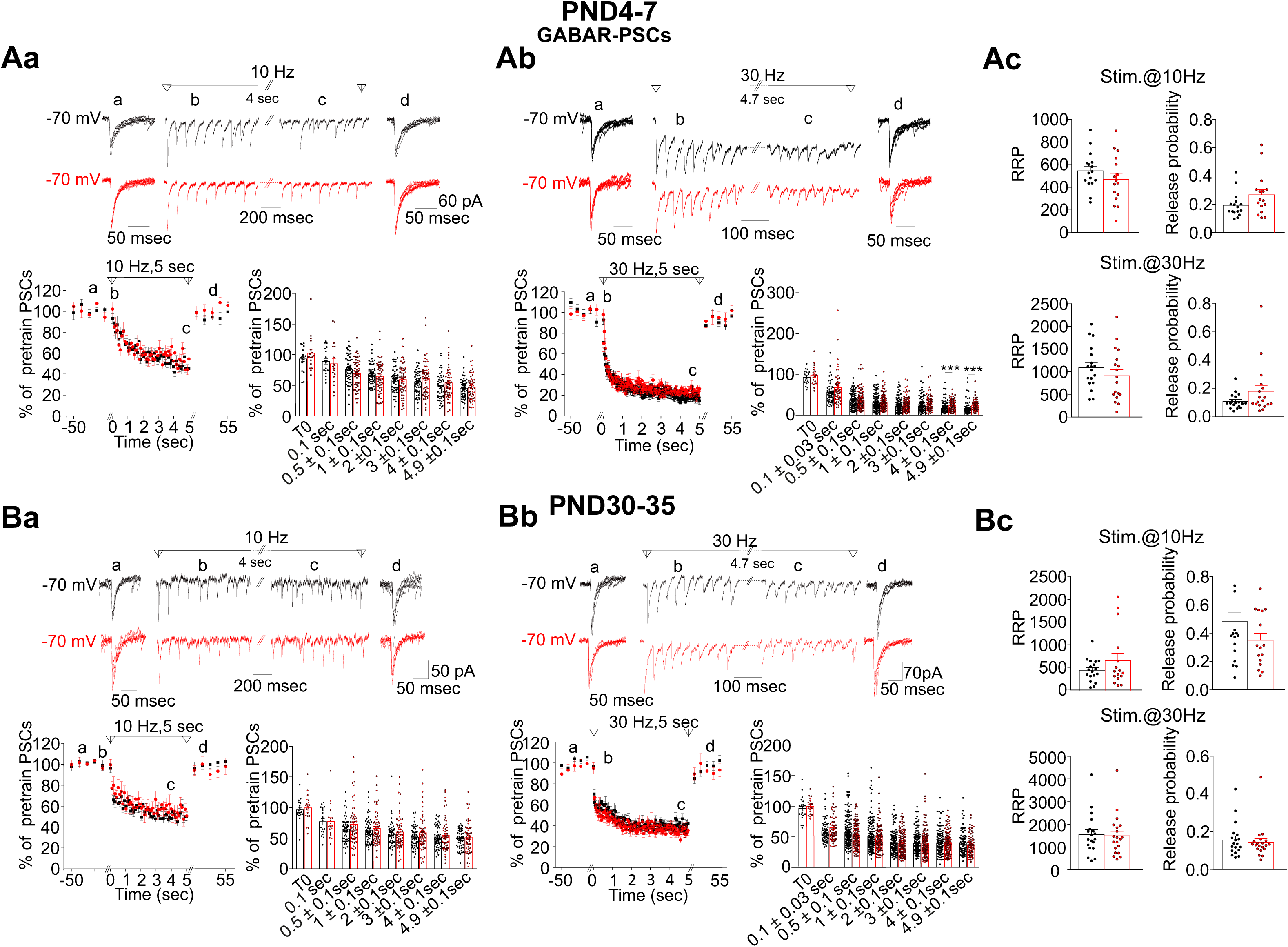
Consequences of Munc18.1 deficiency on GABA receptors mediated postsynaptic currents (GABAR-PSCs) in CA1 pyramidal cells and evoked by electrical stimulation applied at 10 and 30 Hz in the stratum radiatum. **Aa)** GABAR mediated postsynaptic currents (GABAR-PSCs) recorded in CA1 pyramidal cells from WT (black) and *STXBP1^+/−^* mice (red) aged 5 days old. (a) 5 superimposed PSCs evoked at 0.1Hz before the stimulation at 10Hz; (b,c) PSCs recorded during the first and the last second of the stimulation at 10 Hz stimulation; (d) 5 superimposed PSCs evoked at 0.1Hz after the stimulation at 10Hz. Below on the left: Graph quantifying, at PND 4-7, the amplitude of GABAR-PSCs (mean ± SEM, expressed as the percentage of 5 PSCs evoked at 0.1Hz before the stimulation at 10 Hz plus the first PSC of the train). Black: n = 18 cells / 3WT mice in which the stimulation protocol was applied 49 times; Red: n = 18 cells, /3 *STXBP1^+/−^* mice in which the stimulation protocol was applied 54 times. On the right: Boxplots showing average values in each cells of relative GABAR-PSCs during the stimulation at 10 Hz and grouped at times indicated in abscissa. **Ab)** Same as Ba but for stimulations applied at 30Hz during 5 sec. Black: n = 18 cells / 3WT mice in which the stimulation protocol was applied 41 times; Red: n = 18 cells, /3 *STXBP1^+/−^* mice in which the stimulation protocol was applied 53 times. Statistics at t = 4 ± 0.1 sec:*** p <0.001 Mann-Whitney test n = 108 WT GABAR-PSCs, mean ± SEM = 16.8 ± 1.3 % (% of PSCs before the stimulation at 30 Hz) and n = 126 *STXBP1^+/−^* GABAR-PSCs, mean ± SEM = 23.9 ± 1.03 %. At t = 4.9 ± 0.1 sec: *** p <0.001 Mann-Whitney test n = 72 WT GABAR-PSCs, mean ± SEM = 16.6 ± 1.9 % and n = 72 STXBP1^+/−^GABAR-PSCs, mean ± SEM = 24.2 ± 1.5 %. **Ac**) RRP and probability of GABA release from gabaergic inputs located in the stratum radiatum of CA1 from neonatal mice after stimulation at 10 Hz (n = 16 cells/ 3 WT mice and n = 16 cells/3 *STXBP1^+/−^*mice) and at 30 Hz (n = 17 cells/3 WT mice and n = 18 cells/3 *STXBP1^+/−^* mice). **Ba,b**) same as Ba but GABAR-PSCs were recorded in pyramidal cells from juvenile mice aged 30-35 days old. **a)** GABAR-PSC evoked in pyramidal cells by stimulation at 10 Hz during 5 sec. Black: n = 18 cells/4 WT mice in which the stimulation protocol was applied 45 times. Red: n = 19 cells/ 4 *STXBP1^+/−^* mice in which the stimulation protocol was applied 51 times; **b)** GABAR-PSC evoked in pyramidal cells by stimulation at 30 Hz applied during 5 sec. Black: n = 21 cells/3 WT mice in which the stimulation protocol was applied 61 times. Red: n = 20 cells/ 3 *STXBP1^+/−^* mice in which the stimulation protocol was applied 59 times. **Bc**) RRP and probability of GABA release after stimulation at 10 Hz (n = 18 cells/ 4 WT mice and n = 16 cells/4 *STXBP1^+/−^*mice) and at 30 Hz (n = 19 cells/3 WT mice and n = 20 cells/3 *STXBP1^+/−^*mice).

**Figure 5:**
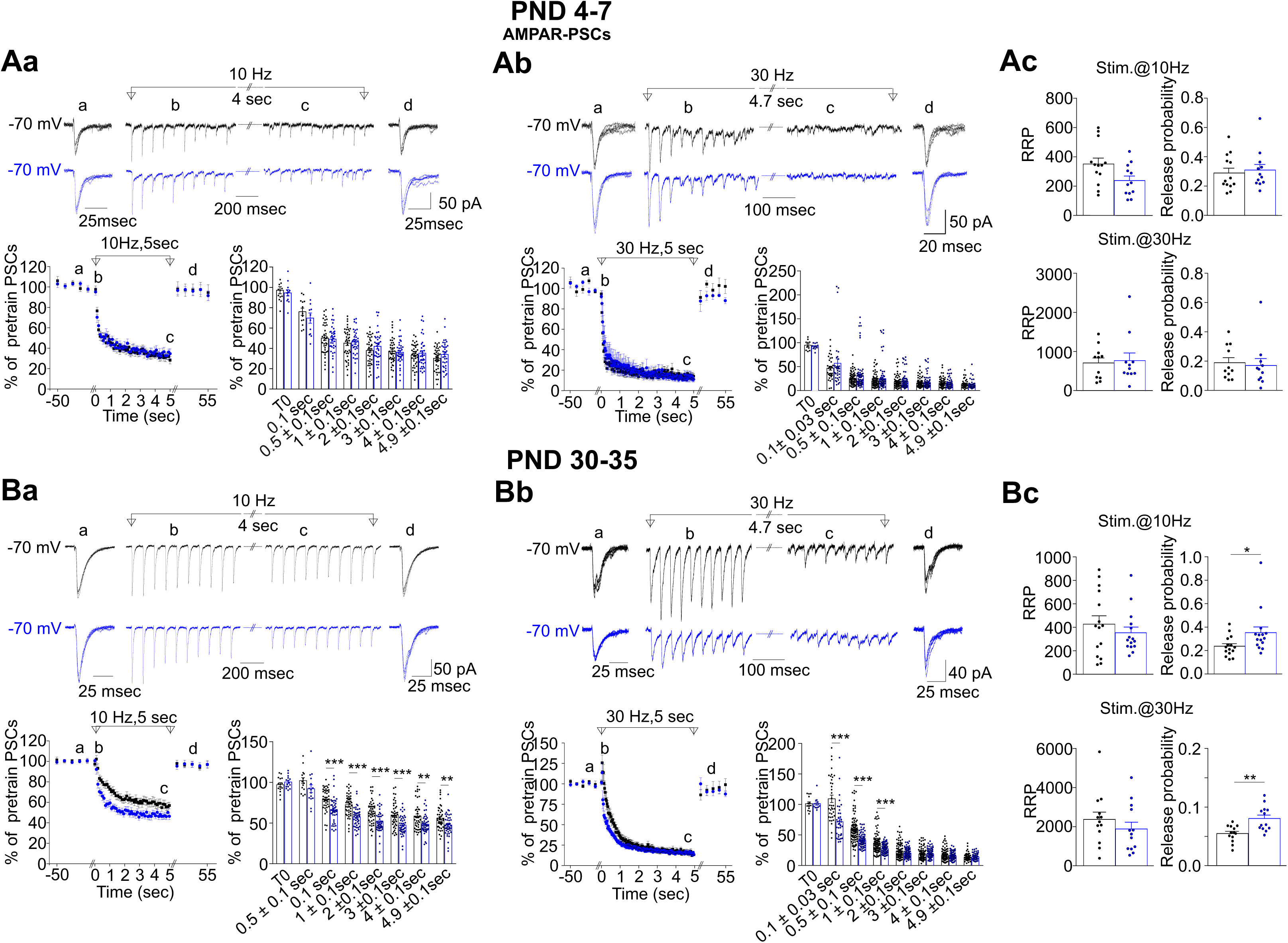
Consequences of Munc18.1 deficiency on AMPAR-PSCs recorded in pyramidal cells of layers II/III of the motor cortex and evoked by electrical stimulation applied at 10 and 30 Hz in the layers II/III. **Aa**) AMPAR mediated postsynaptic currents (AMPAR-PSCs) recorded in Layers II/III pyramidal cells from WT (black) and *STXBP1^+/−^* mice (blue) aged 7 days old. Top: (a) 5 superimposed PSCs evoked at 0.1Hz before the stimulation at 10Hz; (b,c) PSCs recorded during the first and the last second of the stimulation at 10 Hz stimulation (applied during 5 sec); (d) 5 superimposed PSCs evoked at 0.1Hz after the stimulation at 10Hz. Below on the left: Graph quantifying, at PND 4-7, the amplitude of PSCs (mean ± SEM), expressed as the percentage of 5 PSCs evoked at 0.1Hz before the stimulation at 10 Hz plus the first PSC of the train. Black: n = 13 cells / 3WT mice in which the stimulation protocol was applied 30 times; blue: n = 12 cells, /4 *STXBP1^+/−^* mice in which the stimulation protocol was applied 30 times. On the right: Boxplots showing average values in each cells of relative AMPA-PSCs during the stimulation at 10 Hz and grouped at times indicated in abscissa. **Ab)** Same as Aa but for stimulation applied at 30Hz during 5 sec. Black: n = 11 cells / 3WT mice in which the stimulation protocol was applied 21 times; blue: n = 11 cells, /3 *STXBP1^+/−^*mice in which the stimulation protocol was applied 30 times. **Ac)** RRP in layers II/III glutamatergic fibers in motor cortical slices and probability of glutamate release from neonatal WT and *STXBP1^+/−^* mice estimated at 10Hz (Black: n= 13 cells/ 3 WT mice; blue: n = 12 cells/4 *STXBP1^+/−^* mice) and at 30 Hz (Black: n = 11 cells / 3WT mice; blue: n = 11 cells/3 *STXBP1^+/−^* mice). **Ba)** Same as Aa but for AMPAR-PSCs evoked by stimulation at 10 Hz in pyramidal cells from juvenile mice aged 30-35 days old. Traces of from mice aged 30 days old. Black: n = 15 cells/4 WT mice in which the stimulating protocol was applied 39 times; blue: n = 15 cells/3 *STXBP1^+/−^* mice in which the protocol was applied 41 times. Statistics at t = 0.5 ± 0.1 sec: ***p <0.0001 Mann-Whitney test, n = 45 WT AMPAR-PSCs, mean ± SEM = 79.6 ± 2 % and n = 45 *STXBP1^+/−^* AMPAR-PSCs, mean ± SEM = 66.4 ± 2.2 %. Statistics at t = 1 ± 0.1 sec ***p <0.0001 Student t-test, n = 45 WT AMPAR-PSCs, mean ± SEM = 72.6 ± 2 % and n = 45 *STXBP1^+/−^* AMPAR-PSCs, mean ± SEM = 57.3 ± 1.7 %. Statistics at t = 2 ± 0.1 sec: *** p < 0.0001 Mann-Whitney test, n = 45 WT AMPAR-PSCs, mean ± SEM = 64.1 ± 2.2 % and n = 45 *STXBP1^+/−^* AMPAR-PSCs, mean ± SEM = 51.1 ± 1.9 %. Statistics at t = 3 ± 0.1 sec: *** p < 0.0001 Student t-test, n = 45 WT AMPAR-PSCs, mean ± SEM = 59.8 ± 2.2 % and n = 45 *STXBP1^+/−^* AMPAR-PSCs, mean ± SEM = 47.4 ± 1.7 %. Statistics at t = 4 ± 0.1 sec: ** p < 0.0034 Mann-Whitney test, n = 45 WT AMPAR-PSCs, mean ± SEM = 58.7 ± 2.4 % and n = 45 *STXBP1^+/−^*AMPAR-PSCs, mean ± SEM = 48.6 ± 1.7 %. Statistics at t = 5 ± 0.1 sec: ** p < 0.0034 Student t-test, n = 45 WT AMPAR-PSCs, mean ± SEM = 56.7 ± 2.0 % and n = 45 *STXBP1^+/−^* AMPAR-PSCs, mean ± SEM = 47.5 ± 1.9 %. **Bb)** Same as Ba but for stimulations applied at 30 Hz during 5 sec. Black: n = 13 cells/ 3 WT mice in which the stimulation protocol was applied 30 times; Blue: n = 13 cells / 3 *STXBP1^+/−^* mice in which the stimulation protocol was applied 35 times. Statistics at t = 0.1 ± 0.03 sec: ***p <0.0001 Mann-Whitney test, n = 37 WT AMPAR-PSCs, mean ± SEM = 109.3 ± 6.0 % and n = 37 *STXBP1^+/−^* AMPAR-PSCs, mean ± SEM = 71.9 ± 4.9 %. Statistics at t = 0.5 ± 0.1 sec: ***p <0.0001 Mann-Whitney test, n = 91 WT AMPAR-PSCs, mean ± SEM = 59.3 ± 2.2 % and n = 91 *STXBP1^+/−^* AMPAR-PSCs, mean ± SEM = 41.3 ± 1.0 %. Statistics at t = 1 ± 0.1 sec: ***p <0.0001 Mann-Whitney test, n = 91 WT AMPAR-PSCs, mean ± SEM = 36.3 ± 1.7 % and n = 91 *STXBP1^+/−^*AMPAR-PSCs, mean ± SEM = 27.3 ± 0.8 %. **Bc)** RRP and Pr in juvenile mice. At 10 Hz: n = 15 cells/4 WT mice and n = 15 cells/ 3 *STXBP1^+/−^* mice. Statistics probability of glutamate release: *p = 0.019, Mann and Whitney test. Mean ± SEM = 0.24 ± 0.004 in WT and n = 0.35 ± 0.05 in *STXBP1^+/−^*layers II/III glutamatergic fibers. At 30Hz: n = 13 cells/3 WT mice and n = 13 cells/3 *STXBP1^+/−^* mice. Statistics probability of glutamate release: ** p = 0.008, Mann and Whitney test. Mean ± SEM = 0.05 ± 0.004 in WT and n = 0.08 ± 0.007 in *STXBP1^+/−^* layers II/III glutamatergic fibers.

**Figure 6:**
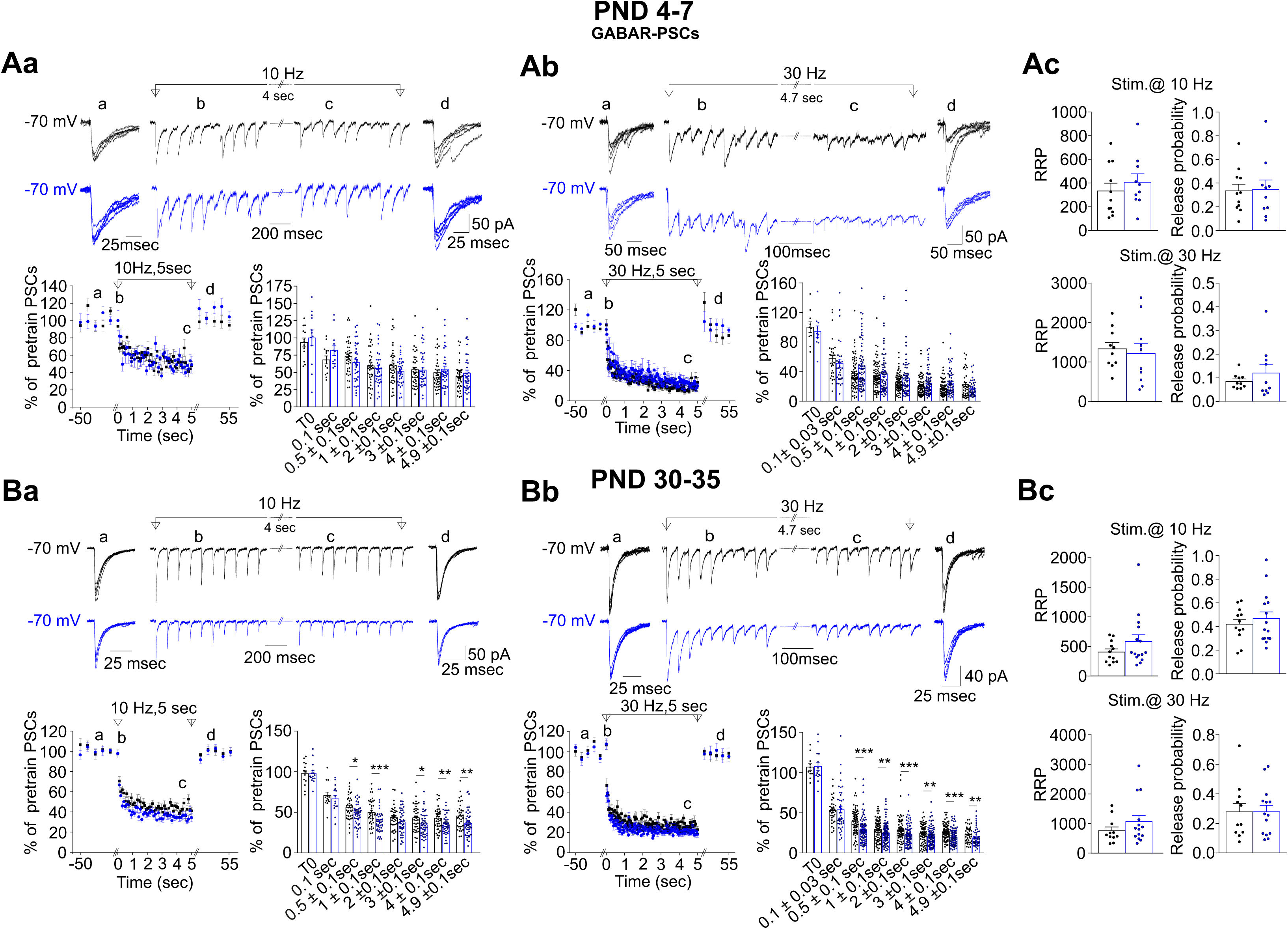
Consequences of Munc18.1 deficiency on GABAR-PSCs recorded in pyramidal cells of layers II/III of the motor cortex and evoked by electrical stimulation applied at 10 and 30 Hz in the layers II/III. **Aa**) GABAR mediated postsynaptic currents (GABAR-PSCs) recorded in pyramidal cells from WT (black) and *STXBP1^+/−^* mice (blue) aged 7 days old. (a) 5 superimposed PSCs evoked at 0.1Hz before the stimulation at 10Hz; (b,c) PSCs recorded during the first and the last second of the stimulation at 10 Hz stimulation; (d) 5 superimposed PSCs evoked at 0.1Hz after the stimulation at 10Hz. Below on the left: Graph quantifying, at PND 4-7, the amplitude of GABAR-PSCs (mean ± SEM), expressed as the percentage of 5 PSCs evoked at 0.1Hz before the stimulation at 10 Hz plus the first PSC of the train. Black: n = 13 cells / 3WT mice in which the stimulation protocol was applied 28 times; blue: n = 12 cells, /3 *STXBP1^+/−^* mice in which the stimulation protocol was applied 32 times. On the right: Boxplots showing average values in each cells of relative GABAR-PSCs during the stimulation at 10 Hz and grouped at times indicated in abscissa. **Ab)** Same as Ba but for stimulations applied at 30Hz during 5 sec (Black: n = 10 cells / 3WT mice in which the stimulation protocol was applied 21 times; blue: n = 10 cells, /3 *STXBP1^+/−^*mice in which the stimulation protocol was applied 30 times. **Ac)** RRP in layers II/III gabaergic fibers in motor cortical slices and probability of GABA release from neonatal WT and *STXBP1^+/−^* mice estimated at 10Hz: n= 11 cells/ 3 WT mice and n = 10 cells/3 *STXBP1^+/−^* mice) and at 30 Hz: n = 10 cells / 3WT mice and n = 10 cells/3 *STXBP1^+/−^* mice. **Ba**) Same as Aa but for GABAR-PSCs evoked by stimulation at 10 Hz and recorded in pyramidal cells from juvenile mice aged 30-35 days old. Black: n = 13 cells / 3WT mice in which the stimulation protocol was applied 36 times; blue: n = 15 cells, /3 *STXBP1^+/−^*mice in which the stimulation protocol was applied 37 times. On the right: Boxplots showing average values in each cells of relative GABAR-PSCs during the stimulation at 10 Hz and grouped at times indicated in abscissa. Statistics at t = 0.5 ± 0.1 sec:** p = 0.0092, Student t-test, n =39 WT GABAR-PSCs, mean ± SEM = 55.8 ± 2.2 % and n = 45 *STXBP1^+/−^* GABAR-PSCs, mean ± SEM = 47.6 ± 2.1 %. Statistics at t = 1 ± 0.1 sec: ***p =0.0004 Mann-Whitney test, n = 39 WT GABAR-PSCs, mean ± SEM = 50 ± 2.5 % and n = 45 *STXBP1^+/−^* GABAR-PSCs, mean ± SEM = 38.5 ± 2.1 %. Statistics at t = 3 ± 0.1 sec: *p = 0.011 Student t-test, n = 39 WT GABAR-PSCs, mean ± SEM = 43.1 ± 2.5 % and n = 45 *STXBP1^+/−^* GABAR-PSCs, mean ± SEM = 34.5 ± 2.1 %. Statistics at t = 4 ± 0.1 sec: **p = 0.004 Student t-test, n = 39 WT GABAR-PSCs, mean ± SEM = 42.7 ± 2.1 % and n = 45 *STXBP1^+/−^*GABAR-PSCs, mean ± SEM = 34.6 ± 1.7 %. Statistics at t = 5 ± 0.1 sec: **p = 0.001 Mann-Whitney test, n = 37 WT GABAR-PSCs, mean ± SEM = 45.4 ± 2.6 % and n = 45 *STXBP1^+/−^*GABAR-PSCs, mean ± SEM = 36.1 ± 2.2 %. **Bb**) Same as Ba but for GABAR-PSCs evoked by stimulation at 30 Hz. Black: n = 11 cells / 3WT mice in which the stimulation protocol was applied 25 times; blue: n = 14 cells, /3 *STXBP1^+/−^* mice in which the stimulation protocol was applied 38 times. Statistics at t = 0.5 ± 0.1 sec: ***p < 0.0001 Mann-Whitney test, n = 77 WT GABAR-PSCs, mean ± SEM = 38.9 ± 1.8 % and n =98 *STXBP1^+/−^* GABAR-PSCs, mean ± SEM = 28.9 ± 1.5 %. Statistics at t = 1 ± 0.1 sec: **p = 0.0065 Mann-Whitney test, n = 77 WT GABAR-PSCs, mean ± SEM = 29.4 ± 1.5 % and n =98 *STXBP1^+/−^*GABAR-PSCs, mean ± SEM = 24.9 ± 1.3 %. Statistics at t = 2 ± 0.1 sec: ***p = 0.0002 Mann-Whitney test, n = 77 WT GABAR-PSCs, mean ± SEM = 28.8 ± 1.8 % and n =98 *STXBP1^+/−^* GABAR-PSCs, mean ± SEM = 21.9 ± 1.2 %. Statistics at t = 3 ± 0.1 sec: **p = 0.0071 Mann-Whitney test, n = 77 WT GABAR-PSCs, mean ± SEM = 24.4 ± 1.3 % and n = 98 *STXBP1^+/−^* GABAR-PSCs, mean ± SEM = 20.6 ± 1.1 %. Statistics at t = 4 ± 0.1 sec: ***p < 0.0001 Mann-Whitney test, n = 77 WT GABAR-PSCs, mean ± SEM = 26.5 ± 1.1 % and n = 98 *STXBP1^+/−^* GABAR-PSCs, mean ± SEM = 19.8 ± 1.0 %. Statistics at t = 5 ± 0.1 sec: **p = 0.0083 Mann-Whitney test, n = 44 WT GABAR-PSCs, mean ± SEM = 23.4 ± 1.4 % and n = 56 *STXBP1^+/−^* GABAR-PSCs, mean ± SEM = 18.8 ± 1.4 %. **Bc)** RRP and probability of GABA release in juvenile. At 10 Hz: n = 12 cells/3WT mice and n = 15 cells/3 *STXBP1^+/−^* mice. At 30 Hz: n = 11 cells/3WT mice and n = 14 cells/3 *STXBP1^+/−^* mice.

Therefore, in both cortical structures, the deficit of Munc18.1 affects the response of the synapses to high frequency stimulation at juvenile but not neonatal stage of development. Moreover, in CA1 as well as in the layers II/III of the motor cortex, glutamatergic synapses appear to be more sensitive than GABAergic synapses to this deficit. This is accompanied by a change in the Pr of glutamate.

### Spontaneous GABA and GluR-PSCs are not altered in neonatal and juvenile heterozygous mice

To complement our study on the impact of Munc18.1 deficiency on synaptic transmission, we analyzed spontaneous synaptic events (in absence of tetrodotoxin), mediated by glutamate and GABA receptors (sGluR-PSCs and sGABAR-PSCs). We compared in WT and *STXBP1^+/−^* pyramidal cells the frequency of these events, their mean amplitude and their amplitude distribution.

At PND4-7 and PND 30-35, spontaneous activity recorded in CA1 was dominated by sGABAR-PSCs (Fig.7). The frequency of which were about 10 times greater than that of sGluR-PSCs (Fig. 7Aa-c, Ba,b). The frequency of sGABAR– and sGluR-PSCs was not significantly different in pyramidal cells recorded from WT and *STXBP1^+/−^* mice at both developmental stages (Fig.7Aa-c, Ba,b). Furthermore, the mean amplitude and amplitude distribution of these spontaneous events were largely insensitive to Munc18.1 deficiency. As already described spontaneous activity recorded in neonatal period may include recurrent outward current (at 0 mV) of few hundreds of pA lasting between 0.5 and 2 s and resulting from the periodic summation of GABAergic-PSC, the so called giant depolarizing potentials (GDPs) in the hippocampus (Ben-Ari et al., 1989). GDPs were recorded in 8 cells in 4 slices from 3 out of 5 WT mice and in 10 cells in 3 slices from 3 out of 6 *STXBP1^+/−^* mice. The frequency and the area of GDPs (corresponding to the charge transfer) measured at 0 mV were not significantly different in WT and *STXBP1^+/−^* pyramidal cells (Fig. 7Ad).

**Figure 7:**
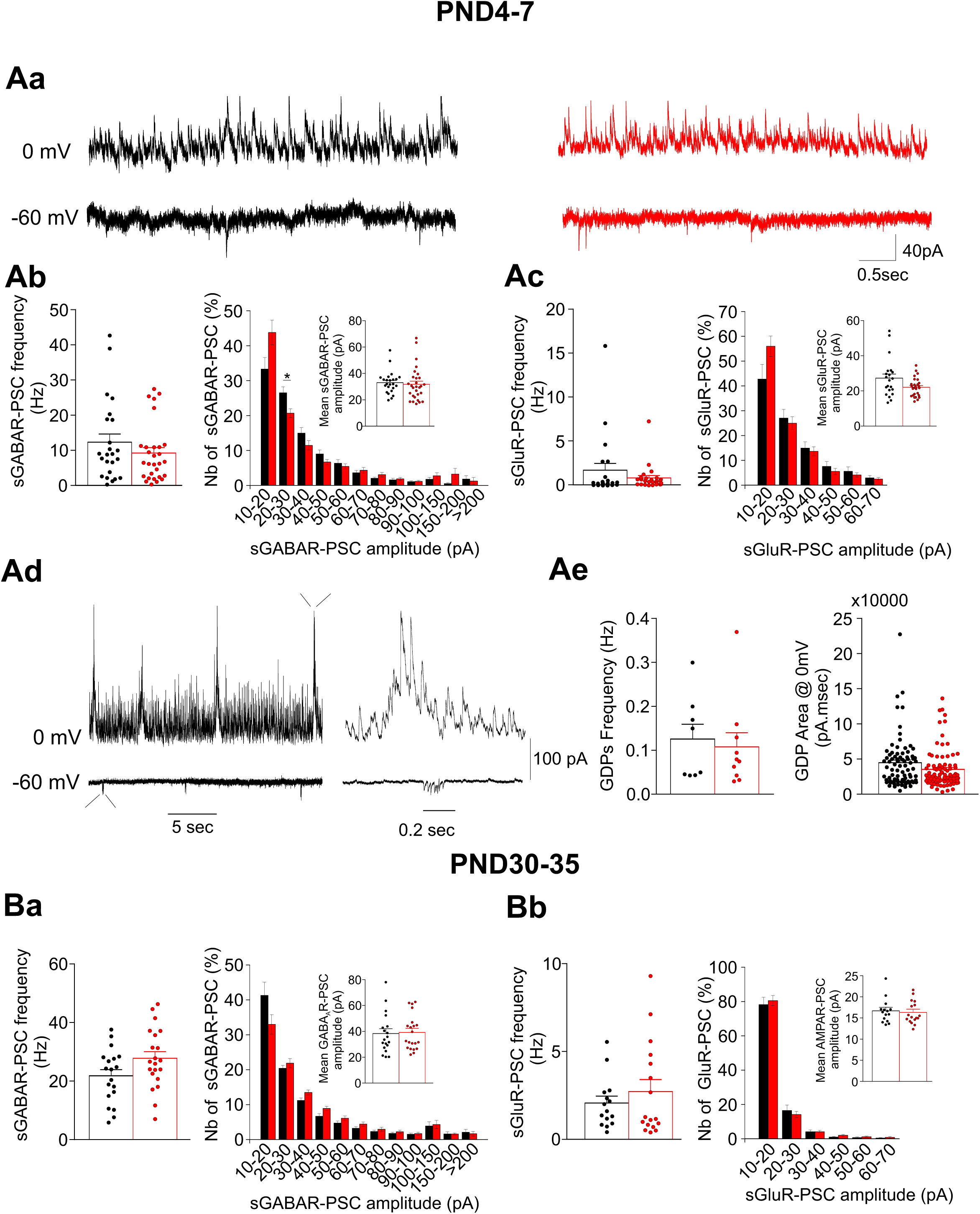
Consequences of a Munc18.1 deficit on the spontaneous ongoing GABAergic and glutamatergic activities in the CA1 region of the hippocampus. **Aa)** Representative traces at PND 6, showing spontaneous activities recorded at 0 and −60 mV in WT and STXBP1^+/−^ CA1 pyramidal cells (black and red traces, respectively). Postsynaptic currents mediated by GABA receptors (sGABAR-PSCs) were recorded at 0 mV, whereas postsynaptic currents mediated glutamate receptors (sGluR-PSCs) were recorded at −60 mV. **Ab,c)** Below the traces are boxplots quantifying at PND4-7 the frequency of sGABAR-PSCs (**b**), sGLuR-PSCs (**c**) and amplitude distribution of the sPSCs. Events were distributed in bin of 10 pA from 10 to 100 pA and in bin of 50 pA for events > 100 pA. Mean amplitude of sGABA and sGluR-PSCs are shown in inserts (sGABAR-PSCs: n= 24 cells/5 WT mice and n = 28 cells/6 *STXBP1^+/−^* mice; sGluR-PSCs: n = 22 cells /5 WT mice and n =27 cells/ 6 *STXBP1^+/−^* mice). **Ad)** Recording of spontaneous activity during 30 sec showing the presence of recurrent large outward currents resulting from the periodic summation of GABAergic-PSCs and underlying giant depolarizing potentials (GDPs) (on enlarged time scale at right). **Ae)** Boxplots quantifying the frequency and the area of the GDPs (calculated on 10 GDPs per cell: n = 8 cells /3 WT mice and n= 10 cells/ 3 *STXBP1^+/−^*mice). **Ba,b)** Boxplots quantifying the frequency of sGABAR-PSCs (**a**), sGLuR-PSCs (**b**) and amplitude distribution of the sPSCs recorded in pyramidal cells from juvenile mice. sGABAR-PSCs: n = 19 cells/3 WT mice and n = 21 cells/ 3 *STXBP1^+/−^* mice. sGluR-PSCs: n = 15 cells/ 3 WT mice and n = 17 cells/ 3 *STXBP1^+/−^*mice.

As in the CA1 region, the spontaneous activity recorded in pyramidal cells of the layers II/III at neonatal and juvenile stages was also mainly mediated by GABAR which frequency was ∼10 times greater than that of sGluR-PSCs (Fig.8A,B). No significant differences were observed in the frequency, mean amplitude and amplitude distribution sGABAR and sGluR-PSCs between neonatal WT and *STXBP1^+/−^* pyramidal cells (Fig.8A,B). In juvenile, the deficit of Munc18.1 did also affect neither the frequency of sGABAR or sGluR-PSCs as well as the amplitude of sGABA events (Fig.8B). A small but significant decrease in the mean amplitude of sGluR-PSCs (on average by 4 pA) was observed in *STXBP1^+/−^*pyramidal cells compared to WT cells probably due to the reduced number of events with amplitudes ranging between 30 and 40 pA (Fig.8Bb).

**Figure 8:**
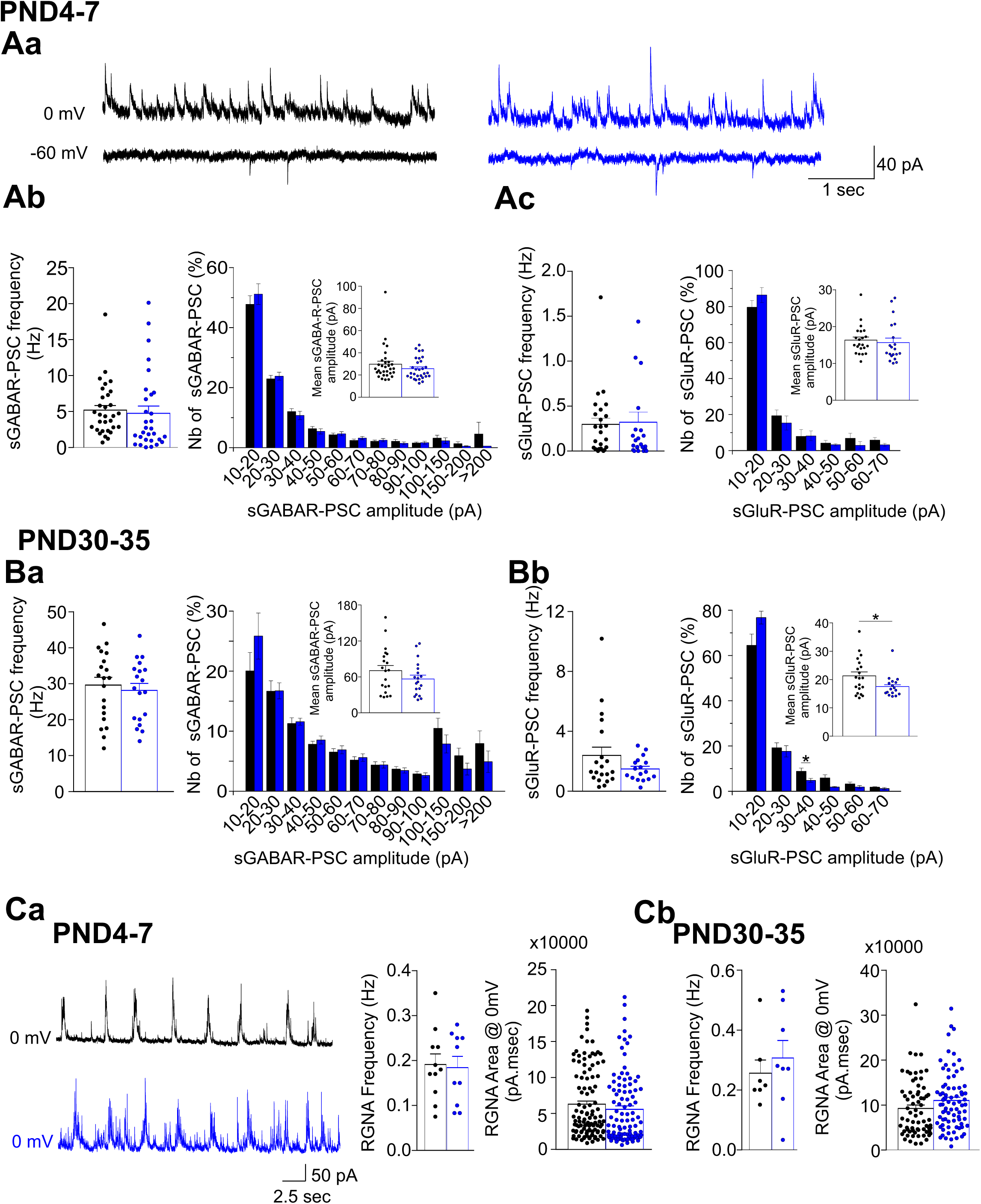
Consequences of a Munc18.1 deficit on the ongoing spontaneous GABAergic and glutamatergic activities in the layers II/III of the motor cortex. **Aa)** Representative traces at postnatal day PND 7, showing spontaneous activities recorded at 0 and −60 mV in WT and STXBP1^+/−^ layers II/III pyramidal cells (black and blue traces, respectively) **Ab,c)** Below the traces are boxplots quantifying at PND4-7 the frequency of sGABAR-PSCs (**b**), sGLuR-PSCs (**c**) and amplitude distribution of the sPSCs. Events were distributed in bin of 10 pA from 10 to 100 pA and in bin of 50 pA for events > 100 pA. Mean amplitude of sGABA and sGluR-PSCs are shown in inserts (sGABAR-PSCs: n= 31cells/4 WT mice and n = 30 cells/4 *STXBP1^+/−^* mice; sGluR-PSCs: n = 22 cells /4 WT mice and n =19 cells/ 4 *STXBP1^+/−^* mice). **Ba,b)** Boxplots quantifying the frequency of sGABAR-PSCs (**a**), sGLuR-PSCs (**b**) and amplitude distribution of the sPSCs recorded in pyramidal cells from juvenile mice. sGABAR-PSC: n = 20 cells/3 WT mice and n = 20 cells/ 3 *STXBP1^+/−^* mice. sGluR-PSC: n = 20 cells/3 WT mice and n = 17 cells/ 3 *STXBP1^+/−^* mice. Statistics amplitude distribution bin 30-40 pA: * p = 0.024, Mann-Whitney test, n = 19WT sGluR-PSCs, mean ± SEM = 8.5 ± 1.4 % and n =16 *STXBP1^+/−^*sGluR-PSCs, mean ± SEM = 4.5 ± 0.9 %. Statistics for mean sGluR-PSC amplitude: *p = 0.011, Student t-test: WT sGluR-PSCs, mean ± SEM = 21.2 ± 1.4 pA; *STXBP1^+/−^*sGluR-PSC, mean ± SEM = 17.1 ± 0.6 pA. **Ca,b)** Recording of Recurrent Gabaergic Network Activity (RGNA) in pyramidal cells from WT and *STXBP1^+/−^*mice aged 5 days old and boxplots quantifying the frequency and the area of the RGNA (calculated on 10 RGNA per cell) at PND 4-7 n = 11 cells / 3 WT mice and n = 10 cells/ 4 *STXBP1^+/−^* mice and PND 30-35: n = 7 cells / 3 WT mice and n = 8 cells/ 2 *STXBP1^+/−^*mice.

GDP like events, which in the motor cortex were named RGNA (recurrent gabaergic network activity, cf Biba-Maazou et al., 2022), were recorded in pyramidal cells not only at PND4-7 but also at PND 30-35 (Fig. 8C). At PND 4-7 they were recorded in 11 cells in 6 slices from 3 out of 3 WT mice and in 10 cells in 6 slices from 4 out of 4 STXBP1^+/−^mice. At PND30-35, RGNA were recorded in 7 cells in 3 slices from 3 out of 3 WT mice and in 8 cells in 4 slices from 2 out of 3 STXBP1^+/−^mice. Neither the frequency nor the area of RGNA was significantly different in WT and STXBP1^+/−^cells at any stage of development.

Therefore Munc18.1 deficiency has no major consequences on spontaneous synaptic activity at the neonatal and juvenile stages of development.

### Target SNARE proteins expression is decreased in juvenile but not in neonatal STXBP1^+/−^ mice

These unexpected time dependent consequences of Munc18.1 deficiency on neuronal excitability and synaptic transmission in neonatal and juvenile mice led us to examine whether these differential electrophysiological consequences were associated with altered expression levels of target SNARE (t-SNARE) proteins at these two developmental stages. Indeed these proteins are involved not only in vesicular exocytosis but t-SNARE proteins modulate the activity of several ionic channels that can potentially affect neuronal excitability (Spafford et al., 2003; Leung et al., 2007). We performed a western blot analysis of Munc18.1, syntaxin 1A, 1B (Stx1A, Stx 1B) and SNAP-25 in the hippocampus and the neocortex from WT and *STXBP1* heterozygous mice at PND 4 and PND30. At PND 4, in the hippocampus of WT and *STXBP1^+/−^* mice, the expression levels of Stx1B and SNAP 25 are higher, while the expression of Munc18.1 and Syx1A are not significantly different (Fig. 9Ab,c). Moreover, as expected, the relative expression of Munc18.1 in the hippocampus of *STXBP1^+/−^* mice is decreased (on average by about 40%) compared to its level in WT mice, while t-SNARE proteins were not affected, leading to Stx1 (1A and 1B)/Munc18.1 and t-SNARE/Munc18.1 ratios that were significantly increased in *STXBP1^+/−^*mice (Fig. 9Aa-d). At PND 30, the relative expression levels of Munc18.1, Stx1A, and SNAP25 remained unchanged in wild-type and *STXBP1^+/−^* mice compared to their respective levels at PND 4 while the level of Stx1B was decreased (Fig. 9C). However, the expression of all these proteins was significantly reduced in *STXBP1^+/−^* mice compared to their expression in wild-type mice (Fig.9B). This led to a similar Syx1/Munc18.1 and t-SNARE/Munc18 ratios in wild-type and *STXBP1^+/−^*mice (Fig. 9Bd).

**Figure 9:**
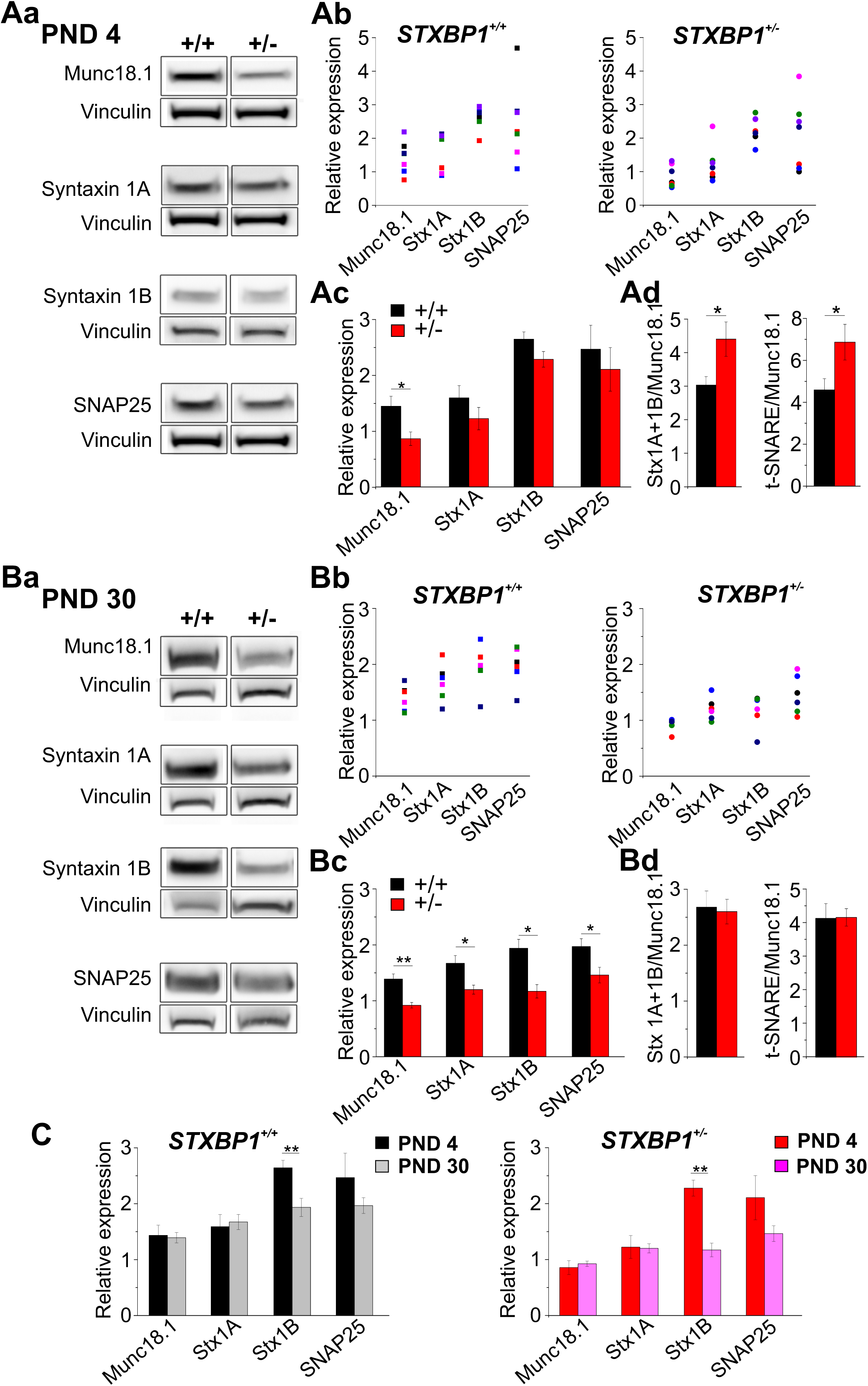
Munc18.1 and t-SNARE proteins expression in the hippocampus from neonatal and juvenile *STXBP1* heterozygous mice. **Aa**) Western blots of Munc18.1, Syntaxin 1A (Stx 1A), Syntaxin 1B (Stx 1B), SNAP25 and Vinculin at postnatal day 4 (PND 4) in the whole hippocampus from WT (+/+) and STXBP1 heterozygous (+/−) mice. **Ab,c**) Quantification of Munc18.1 and t-SNARE proteins expression relative to the housekeeping protein Vinculin in the two mouse groups. Each colored dot/square corresponds to quantification in one mouse. The mean values (± SEM) for each protein are shown in Ac in histograms (black: WT mice; red: STXBP1^+/−^ mice). Statistics Munc18.1: * p = 0.026 Mann and Whitney test, n = 7 WT mice, mean ± SEM = 1.43 ± 0.18 and n = 7 *STXBP1^+/−^* mice, mean ± SEM = 0.85 ± 0.12. **Ad)** Histograms showing the calculation (mean ± SEM) of the Stx 1 (1A + 1B)/ Munc18.1 and t-SNARE/ Munc18.1 ratios (left and right histograms respectively) in WT and *STXBP1^+/−^*mice. Statistics for Stx1/Munc18.1 ratio: * p = 0.025 Mann-Whitney test, n = 7 WT mice, mean ± SEM = 3.0 ± 0.2 and n = 7 *STXBP1^+/−^* mice, mean ± SEM = 4.4 ± 0.5. Statistics for t-SNARE/Munc18.1 ratio: * p = 0.026 Mann-Whitney test, n = 7 WT mice, mean ± SEM = 4.6 ± 0.5 and n = 7 *STXBP1^+/−^*mice, mean ± SEM = 6.9 ± 0.8. **B)** Same analysis than in A but at PND 30. Statistics for Munc18.1 relative expression: ** p = 0.002 Mann and Whitney test, n = 6 WT mice, mean ± SEM = 1.4 ± 0.09 and n = 6 *STXBP1^+/−^* mice, mean ± SEM = 0.9 ± 0.05. Statistics for Stx1A relative expression * p = 0.02 Mann and Whitney test, n = 6 WT mice,, mean ± SEM = 1.7 ± 0.1 and n = 6 *STXBP1^+/−^*mice, mean ± SEM = 1.2 ± 0.08. Statistics for Stx1B relative expression * p = 0.015 Mann and Whitney test, n = 6 WT mice,, mean ± SEM = 1.9 ± 0.1 and n = 6 *STXBP1^+/−^* mice, mean ± SEM = 1.2 ± 0.1. Statistics for SNAP25 relative expression * p = 0.02 Mann and Whitney test, n = 6 WT mice, mean ± SEM = 2 ± 0.1 and n = 6 *STXBP1^+/−^* mice, mean ± SEM = 1.4 ± 0.1. **C)** Histograms showing the evolution of Munc18.1 and t-SNARE proteins relative expressions at PND 4 and PND 30 in the hippocampus of WT (left histograms) and *STXBP1^+/−^*mice (right histograms). Statistics for Stx1B relative expression in WT mice: ** p = 0.001 Mann and Whitney test, n = 7 mice at PND 4, mean ± SEM = 2.6 ± 0.1 and n = 6 mice at PND 30, mean ± SEM = 1.9 ± 0.2. Statistics for Stx1B relative expression in STXBP1^+/−^ mice: ** p = 0.001 Mann and Whitney test, n = 7 mice at PND 4, mean ± SEM = 2.3 ± 0.1 and n = 6 mice at PND 30, mean ± SEM = 1.2 ± 0.1.

In the neocortex at PND 4, a difference is observed with the hippocampus, particularly concerning the SNAP25 protein, whose relative expression level in wild-type and heterozygous mice was lower than that of other proteins (Fig. 10A). However, as in the hippocampus, only the relative expression of Munc18.1 was reduced (by about 50%) in *STXBP1^+/−^* mice compared to its expression level in WT mice, resulting in a significant increase in Syx1/Munc18.1 and t-SNARE/Munc18.1 ratios (Fig. 10Ad). At PND 30, the relative expression of the proteins evolves similarly in WT and *STXBP1^+/−^*mice compared to their respective levels at PND 4, with an increase in the levels of Stx1A and SNAP25 (and Stx1B only in WT) (Fig 10C). However, as in the hippocampus, the expression levels of t-SNARE proteins, as well as Munc18.1, were reduced in *STXBP1^+/−^* mice compared to wild-type mice (Fig. 10Ba-c). The calculation of Stx1/Munc18.1 and t-SNARE/Munc18.1 ratios no longer showed a significant difference between the two groups of mice (Fig. 10Bd).

**Figure 10:**
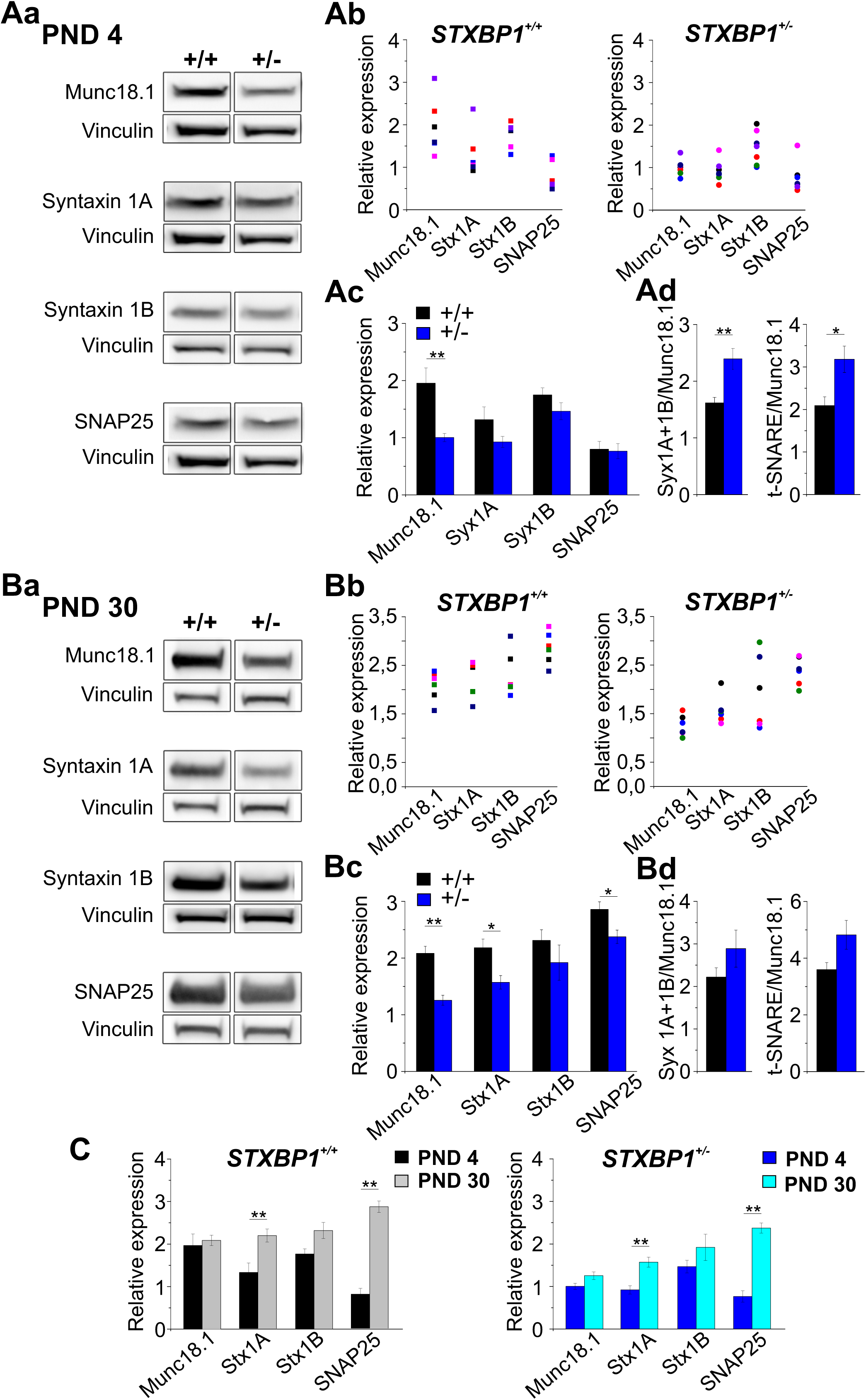
Munc18.1 and t-SNARE proteins expression in the neocortex from neonatal and juvenile *STXBP1* heterozygous mice. **Aa**) Western blots of Munc18.1, Syntaxin 1A (Stx 1A), Syntaxin 1B (Stx 1B), SNAP25 and Vinculin at postnatal day 4 (PND 4) in the whole hippocampus from WT (+/+) and STXBP1 heterozygous (+/−) mice. **Ab,c**) Quantification of Munc18.1 and t-SNARE proteins expression relative to the housekeeping protein Vinculin in the two mouse groups. Each colored dot/square corresponds to quantification in one mouse. The mean values (± SEM) for each protein are shown in Ac in histograms (black: WT mice; Blue: *STXBP1^+/−^*mice). Statistics for Munc18.1: ** p = 0.0012 Mann and Whitney test, n = 6 WT mice, mean ± SEM = 2 ± 0.3 and n = 7 *STXBP1^+/−^* mice, mean ± SEM = 1.0 ± 0.07. **Ad)** Histograms showing the calculation (mean ± SEM) of the Stx1 (1A + 1B)/ Munc18.1 and t-SNARE/ Munc18.1 ratios (left and right histograms respectively) in WT and *STXBP1^+/−^* mice. Statistics for Stx1/Munc18.1 ratio: ** p = 0.005 Mann-Whitney test, n = 6 WT mice, mean ± SEM = 1.6 ± 0.1 and n = 7 *STXBP1^+/−^* mice, mean ± SEM = 2.4 ± 0.2. Statistics for t-SNARE/Munc18.1 ratio: * p = 0.02 Mann-Whitney test, n = 6 WT mice, mean ± SEM = 2.1 ± 0.2 and n = 7 *STXBP1^+/−^*mice, mean ± SEM = 3.2 ± 0.3. **B)** Same analysis than in A but at PND 30. **Bc**) Statistics for Munc18.1 relative expression: ** p = 0.004 Mann and Whitney test, n = 6 WT mice, mean ± SEM = 2.1 ± 0.1 and n = 6 *STXBP1^+/−^* mice, mean ± SEM = 1.3 ± 0.09. Statistics for Stx1A relative expression * p = 0.01 Mann and Whitney test, n = 6 WT mice, mean ± SEM = 2.2 ± 0.2 and n = 6 *STXBP1^+/−^*mice, mean ± SEM = 1.6 ± 0.1. Statistics for SNAP25 relative expression * p = 0.04 Mann and Whitney test, n = 6 WT mice, mean ± SEM = 2.9 ± 0.1 and n = 6 *STXBP1^+/−^*mice, mean ± SEM = 2.4 ± 0.1. **C)** Histograms showing the evolution of Munc18.1 and t-SNARE proteins relative expressions at PND 4 and PND 30 in the neocortex of WT (left histograms) and *STXBP1^+/−^*mice (right histograms). Statistics for Stx1A relative expression in WT mice: * p = 0.013 Mann and Whitney test, n = 6 mice at PND 4, mean ± SEM = 1.3 ± 0.2 and n = 6 mice at PND 30, mean ± SEM = 2.2 ± 0.1. Statistics for Stx1B relative expression in WT mice: *p = 0.03 Mann and Whitney test, n = 6 mice at PND 4, mean ± SEM = 1.8 ± 0.1 and n = 6 mice at PND 30, mean ± SEM = 2.3 ± 0.2. Statistics for SNAP25 relative expression in WT mice: **p = 0.0012 Mann and Whitney test, n = 6 mice at PND 4, mean ± SEM = 0.8 ± 0.1 and n = 6 mice at PND 30, mean ± SEM = 2.9 ± 0.1. Statistics for Stx1A relative expression in *STXBP1^+/−^* mice: ** p = 0.005 Mann and Whitney test, n = 6 mice at PND 4, mean ± SEM = 0.9 ± 0.09 and n = 6 mice at PND 30, mean ± SEM = 1.6 ± 0.1. Statistics for SNAP25 relative expression in *STXBP1^+/−^*mice: ** p = 0.0012 Mann and Whitney test, n = 6 mice at PND 4, mean ± SEM = 0.8 ± 0.1 and n = 6 mice at PND 30, mean ± SEM = 2.4 ± 0.1.

These data therefore show that between PND 4 and PND 30 in the hippocampus as well as in the neocortex, expression levels of the Munc18.1 and t-SNARE proteins follows the same evolution in wild type and heterozygous mice, but the level of these proteins is less important in *STXBP1^+/−^* than in WT mice.

## Discussion

In this study, we conducted electrophysiological and biochemical analyses to evaluate the impact of Munc18.1 deficiency on the intrinsic properties of cortical pyramidal cells, as well as on evoked and spontaneous synaptic transmission and on the content of Munc18.1 and t-SNARE proteins in the neocortex and hippocampus during neonatal and juvenile developmental stages. Until now, the electrophysiological analyses performed *ex-vivo* from various Munc18.1 deficient mouse models have been conducted after weaning or in adulthood, but never during the neonatal period (Orock et al., 2018; Miyamoto et al., 2019; Chen et al., 2020; Dos Santos et al., 2023). This is important for understanding pathophysiological mechanisms of developmental epileptic encephalopathies (DEE) associated with Munc18.1 haploinsufficiency because Munc18.1 is expressed already at embryonic stage, controlling neuronal migration (Hamada et al., 2017; but see Verhage et al., 2000) and because the first clinical manifestations of patients with STXBP1 haploinsufficiency are already observed in newborns (Saitsu et al., 2008; Di Meglio et al., 2015; Stamberger et al., 2016; Abramov et al., 2021). Furthermore, a mechanism could be present in the neonatal period, but limited in time and not observed at a later stage of development, and conversely, only manifest after the network has reached a certain stage of development (see Biba-Maazou et al., 2022; Mao et al., 2025; Reva et al., 2025; Wang et al., 2025). This is clearly what the present study seems to indicate. We show here that Munc18.1 deficiency has different electrophysiological consequences in neonates and juvenile mice. In neonates, the cellular intrinsic excitability is affected, while in juvenile mice, the evoked synaptic transmission is impacted. Furthermore, we observed that a Munc18.1 deficit in heterozygous mice is associated with decreased t-SNARE proteins expression levels in juvenile mice, but not in neonates. Therefore, Munc18.1 deficiency has multiple consequences which depend on the stage of development.

Thus, during the neonatal period, we observed that pyramidal cells located in the CA1 region of the hippocampus and in layers II/III of the motor cortex were more excitable in *STXBP1* heterozygous mice than in wild-type mice. Interestingly, Munc18.1 deficiency had same electrophysiological consequences in both populations of pyramidal cells. Their firing rate was increased in response to depolarizing currents steps above 40 pA; this hyperexcitability was associated with an increase in the final but not the initial discharge frequency. Moreover, the amplitude of AP elicited by a short depolarizing step command was also stronger, and no alterations in intrinsic cell properties (Vm, Rm measurement at rest, membrane time constant) were observed. This indicates that these alterations may have the same origin in both structures and suggests that Munc18.1 deficiency leads to a change in the function/properties of the same ion channel(s), which, according to our data, would likely contribute to the adaptation of neuronal discharge and AP amplitude.Therefore, in addition to its role in transmitter release, Munc18.1 could control the activity of ion channels. However, to our knowledge, no interaction of Munc18.1 with any of the K^+^, Na^+^, or Ca^2+^ channels has been reported so far. This is not the case for the Munc18.1 partner syntaxin I (Stx1) or SNAP25, which has been shown to inhibit in heterologous cells the activity of several Kv channels (Kv1, Kv2, Kv4, Kv7) (Fili et al.,2001; Michaelevski et al; 2003; Leung et al.,2007; Regev et al., 2009; Devaux et al., 2017) as well as voltage-gated Ca^2+^ channels, mainly Cav2.1 (P/Q type) and Cav2.2 (N type) (Bezprozvanny et al., 1995; Wiser et al., 1996; Atlas, 2001; Jarvis and Zamponi, 2001,Jarvis et al., 2002; Spafford and Zamponi; 2003, Gladisheva et al. 2004; Condliffe et al., 2010; Pozzi et al., 2019), epithelial Na^+^ channels (Saxena et al., 1999; Peters et al., 2001), cyclic AMP-gated chloride channel (CFTR chloride channel, Naren et al. 1997; Peters et al., 2001), by a direct physical interaction that can affect the gating properties of the channels as well as their trafficking to the membrane. Interestingly, Munc18.1 was found to abrogate the inhibitory impact of Stx1A on several ion channels (Naren et al., 1997; Jarvis and Zamponi, 2001; Gladisheva et al. 2004; Devaux et al., 2017) via a mechanism that prevent the interaction of the t-SNARE proteins with ion channels (Devaux et al., 2017). It is therefore possible that the same mechanisms operate in neurons and that the expression level of Munc18.1 in *STXBP1* heterozygous mice is insufficient to prevent the inhibitory action of t-SNARE proteins on ion channels in particular Kv channels leading to an increase in neuronal discharge and in the amplitude of AP. For example, the full abrogation by Munc18.1 of the inhibition of CFTR chloride channel by Stx1A is observed only after a certain level of Munc18.1 expression is reached (Naren et al., 1997). Interestingly, in the hippocampus and neocortex, we found that the expression of Munc18.1 relative to Stx1 or t-SNARE proteins was significantly decreased at PND 4 in *STXBP1* heterozygous mice compared to wild-type mice, but not different at PND 30, when neuronal excitability is restored. We do not yet have evidence that the relative expression levels of these proteins is decisive for neuronal excitability; this is simply a correlation and we cannot exclude that other mechanisms are responsible for both the hyperexcitability of cells at PND 4 and the normalization of the firing pattern of pyramidal cells and AP amplitude at PND 30. It is noteworthy that an increase of neuronal discharge was not observed in neurons derived from conditional mutant human embryonic stem cells, but it is interesting to note that in these mutant cells, a decrease in Munc18.1 expression was also associated with a decrease in Stx1 expression (Patzke et al. 2015). Clearly, further important biochemical and electrophysiological studies are needed to understand the origin and mechanism involved in this time-limited increase in pyramidal cells hyperexcitability.

At the juvenile stage, we observed that synaptic transmission is affected in *STXBP1* heterozygous mice when the synaptic response was evoked by stimulations at 10 and 30 Hz. However, regarding AMPAR-PSC, we found differences in the sensitivity of responses to stimulus trains in CA1 pyramidal cells and in layers II/III in wild type and *STXBP1* heterozygous mice. In wild type mice, after 10 Hz stimulation, the response was increased in CA1 but decreased in layers II/III while the estimated size of the readily releasable pool of vesicles (RRP) and the probability of release (Pr) were similar in presynaptic fibers stimulated in the 2 regions (see Figs.4C and 6C). Pre– and postsynaptic mechanisms could potentially explain these effects, including a difference in the rate of RRP replenishment, in the composition and properties of presynaptic calcium channels (in particular their inactivation; Forsythe et al. 1998), the composition of postsynaptic AMPA receptors mediated-EPSCs, and their auxiliary proteins that together would determine the level of receptor desensitization (Greger et al., 2017; Kamalova and Nakagawa, 2021). More surprising was the enhancement of the synaptic response at 10 Hz in CA1 of *STXBP1* heterozygous mice at least during the first 3 sec of the train whereas we expected either no effect or a run-down of EPSCs as observed in other *ex-vivo* studies (Miyamoto et al., 2019; Chen et al., 2020; Dos Santos et al., 2023). For example, in Miyamoto’s study, stimulation at 10 Hz of synapses made by pyramidal cells located in layers V/VI of the somatosensory cortex and the fast-spiking interneurons of the striatum led to a facilitation of AMPAR-PSC in wild type mice but to a depression in *STXBP1* heterozygous mice. In our study, a significant larger depression of AMPAR-PSC was however observed at 30Hz in the hippocampus of *STXBP1* heterozygous compared to wild type mice after an initial enhancement of the response. In fact, our data revealed that Munc18.1 deficiency affected Pr but not the RRP in CA1 and in the layers II/III. The lack of consequences of Munc 18.1 deficiency on RRP was however surprising because in both structures decrease level of Munc18.1 was also associated with a decrease in expression level of t-SNARE proteins which together should affect the priming and fusion of synaptic vesicles (Sudhöf 2014; Vardar et al., 2016). One possible explanation is that the overall decrease in these proteins is not severe enough to significantly affect the RRP (Arancillo et al., 2013, but see Toonen et al. 2006). In any case, our data are consistent with other studies performed either in acute brain slices where the synaptic response to trains of stimulation was impacted, and the RRP size estimated (Miyamoto et al., 2019) or in neurons derived from conditional mutant human embryonic stem cells (Patzke et al. 2015). It is also the case in studies in which *STXBP1* heterozygous neurons support normal synaptic transmission, data that were obtained in neuronal culture expressing human STXBP1 variants (Kovacevic et al, 2018) and in *STXBP1* patients-IPSC derived neurons (Van Berkel et al. 2024).

However, to our knowledge, no increase in synaptic response during the train of stimulation or change in Pr has been observed on any *STXBP1* heterozygous synapses analyzed so far *ex-vivo* and *in vitro* (Toonen et al., 2006; Miyamoto et al. 2019; Chen et al., 2020; Dos Santos et al. 2023). The change in Pr suggests some presynaptic regulation at the Ca^2+^ level (dynamics, flow through voltage-gated Ca^2+^ channels, proximity between Ca^2+^ channels and vesicles, sensitivity of vesicles to Ca^2+^…, Dittman and Ryan, 2019) in which the reduction in expression level of t-SNARE proteins could potentially play a role (Bergsman and Tsien 2000; Verderio et al., 2004; Condliffe et al., 2010; Antonucci et al., 2013; Pozzi et al.,2019) but the fact that Pr changed in opposite direction in CA1 and in layers II/III is difficult to explain and requires further investigation.

Thus, the functional consequences of Munc18.1 deficiency in synaptic transmission are not homogeneous for all synapses. This is also supported by our analysis of GABA receptor-mediated synaptic transmission. Stimulation at 10 and 30 Hz induced a rapid and significant decrease in the GABAR-PSC in wild-type mice, both in CA1 and in layers II/III. This decrease was significantly, although slightly stronger, in *STXBP1* heterozygous mice (by 10–15%) in layers II/III, while the depression level was similar in CA1. Therefore, although Munc18.1 plays a fundamental role in neurotransmission, not all synapses are affected equally by protein deficiency even within the same structures, some synapses being sensitive while others insensitive to reduced Munc18.1 expression. In keeping with this, an insensitivity to a deficit of Munc18.1 of glutamate and GABA receptors-mediated synaptic transmission to high frequency stimulation has been reported *ex-vivo* at synapses made by pyramidal cells of layers V/VI of the somatosensory cortex and the medial spiny interneuron of the striatum, whereas the same synapses made on fast spiking interneuron produced a faster and stronger rundown of AMPAR-PSC (Miyamoto et al. 2019). A sensitivity to a deficit of Munc18.1 has been observed in somatosensory cortex at glutamatergic synapses made between layer IV pyramidal cells and layers II/III pyramidal cells or PV positive interneurons while GABAergic synapses made in same layers by PV or SST positive interneurons and pyramidal cells were unaffected, in total creating an imbalance between excitation and inhibition (Chen et al; 2020; Dos Santos et al., 2023). The reason why a Munc18.1 deficit affects transmitter release to train of stimulation in some synapses (mainly glutamatergic synapses) with much less effect on GABAergic synapses at least in *ex-vivo* conditions (see however Toonen et al., 2006 in neuronal culture performed from *STXBP1* heterozygous mice) while Munc18.1 is express in both population of cells remains to be elucidated. However, it is possible that changes in expression levels of Munc18.1 and its partners are different in pyramidal cells and in interneurons of *STXBP1* heterozygous mice. Our biochemical analysis shows global effects in whole neocortex and hippocampus and does not allow us to distinguish the expression of these proteins in the different population of cells. Interestingly, differences exist between pyramidal cells and interneurons regarding t-SNARE proteins content. GABAergic interneurons do not express SNAP25 or at much lower level than glutamatergic neurons, and they evoked larger calcium current to depolarizing stimuli than pyramidal neurons likely due to the absence of SNAP25 which exerts a negative regulation of voltage gated calcium channels (Verderio et al.,2004; Condliffe et al., 2010; Pozzi et al., 2019). Because Ca^2+^ regulates the mobilization, the priming and fusion of synaptic vesicles, a larger calcium influx may perhaps counterbalance the impact that a decrease in Munc18.1 content in presynaptic terminal may have on neurotransmission.

We did not observe major differences in GABAR– and AMPAR-mediated spontaneous activity in layers II/III and in the CA1 region of neonatal or juvenile heterozygous mice compared to wild-type mice indicating that Munc18.1 deficiency does not impact ongoing synaptic activity in cortical networks at any developmental stages. However, we were surprised that at PND 4-7, spontaneous activity mediated by AMPA receptors was not increased in *STXBP1* heterozygous mice, while pyramidal neurons were hyperexcitable. It is possible that this hyperexcitability is not generalized to all neonatal neocortical and hippocampal pyramidal cells. Indeed, the ion channel(s) potentially affected by Munc18.1 deficiency may not develop and be expressed at same time in all pyramidal neurons, or may have no major impact on their excitability. For example, Kv7 channels-mediated M-current develops at an earlier stage in layers II/III than in layer V pyramidal cells, and their blockade during the neonatal period increases the excitability of the former but not of the latter cells (Biba-Maazou et al., 2022). Another possibility is that the variation in the membrane potential of pyramidal neurons under ongoing conditions is not sufficient to reach the level of depolarization where neuronal firing is significantly increased. Indeed, the shift in the relationship between the number of action potentials and the current injected that was observed in *STXBP1* heterozygous cells in comparison to wild type cells is only observed from a certain level of depolarization. The situation is likely to be different *in vivo* where neuromodulator and external sensory cues, in addition to synchronization through different rhythmic activities/oscillatory patterns present in neonatal brain, such as spindle burst, early gamma oscillations and early sharp waves, would favor membrane depolarization and neuronal firing (Khazipov and Milh, 2018; Valeeva et al., 2019).

To our knowledge, only one study has analyzed ongoing spontaneous events (in the absence of TTX) and reached similar conclusions. Thus, it was shown that spontaneous AMPAR-PSC recorded in PV and SST positive interneurons of the somatosensory cortex were not sensitive (frequency and amplitude) to Munc18.1 deficiency (Chen et al. 2020). In other studies it was the miniatures EPSC and IPSC that were analyzed and both an insensitivity to Munc18.1 deficiency or an increase or decrease in the amplitude or frequency of the miniatures have been reported (Toonen et al., 2006; Patzke et al., 2015; Kovacevic et al., 2020; Dos Santos et al, 2023, McLeod et al., 2023; Van Berkel et al., 2024)

The main finding of our biochemical analysis is the decrease in the relative expression levels of t-SNARE proteins in STXBP1 heterozygous juvenile mice, but not in neonatal mice. This latter observation is consistent with western blot analysis of these proteins in the whole brain of the same STXBP1 heterozygous mouse model at the late embryonic stage (Toonen et al., 2005). We observed both in the hippocampus and neocortex that the expression profile of the different proteins is the same in WT and heterozygous mice, at PND4 or PND 30, but the evolution of t-SNARE expression from PND 4 to PND 30 occurs at a lower level in *STXBP1* heterozygous mice compared to wild type mice. A decrease in expression of Stx1 has been also reported in neurons derived from heterozygous *STXBP1* mutant human ES cells (Patzke et al., 2015), in neurons derived from *STXBP1* patients while RNA levels were unaffected, suggesting posttranslational modification (Van Berkel et al., 2024) and in adult brains of *STXBP1* heterozygous mice (Guiberson et al., 2024). An important question is why the reduction of t-SNARE protein expression does not occur at PND 4 but at PND 30. It is known that Munc18.1 and Stx1, at least, are chaperones to each other, the deficiency of Munc18.1 leading to the instability of Stx1 and vice versa (Toonen et al., 2005, Arancillo et al., 2013; Guiberson et al., 2024) or the increase expression of Munc18.1 increases the expression of Stx1 and of the other chaperone Doc2A/B (Guiberson et al., 2024). It may be that the decrease in Munc18.1 expression level is not strong enough to affect expression of t-SNARE proteins at PND 4. Indeed, decreased levels of Stx1 and Doc2A/B were observed in whole brain of *STXBP1* homozygous where Munc18.1 was absent, but not heterozygous mice (Toonen et al., 2005) although other studies observed reduction of these two chaperones proteins in heterozygous condition where Munc18.1 level was decreased by only 20 to 50% (Patzke et al., 2015; Guiberson et al., 2024;Van Berkel et al., 2024). Here it seems unlikely that the reduced level of t-SNARE proteins observed at PND 30 but not at PND 4 is related to the decrease of Munc18.1 since there was no difference in the relative expression of Munc18.1 in *STXBP1* heterozygous mice at these two developmental stages. Furthermore SNAP 25 which does not directly interact with Munc18.1 was also decreased at PND 30 and it has been shown that the expression of this protein is not sensitive to Munc18.1 expression level (Toonen et al., 2005; Patzke et al., 2015). Altogether, we suggest that the downregulation of the t-SNARE proteins in *STXBP1* heterozygous mice brain observed in our study is the consequence of modification of neuronal activity that the deficit of Munc18.1 produces. With this hypothesis, it would be interesting to see how decreasing levels of Stx1 and SNAP25 at PND 4 or increasing their expression at PND 30 affects the firing rate of pyramidal neurons and synaptic transmission evoked by high frequency stimulation.

In conclusion, the potential impact of Munc18.1 deficiency on the intrinsic properties and firing pattern of neurons has not been fully investigated. Our data suggest that a dysfunction of ionic channels leading to a transient neuronal hyperexcitability must be integrated into the pathophysiology of *STXBP1* related disorders. Our data suggest that it is in fact the first electrophysiological consequences of a deficit in Munc18.1 expression. Interestingly, a time-limited increase in neuronal excitability has also been demonstrated in mice model of *KCNQ2* and *SCNA2-*mediated DEE (Biba-Maazou et al., 2022; Wang et al; 2025; Reva et al., 2025). It therefore appears that early and transient neuronal hyperexcitability is a characteristic of DEE whatever its origin and that a compensatory mechanism develops with time to restore a normal firing pattern of pyramidal neurons at least. We suggest here, without providing evidence yet, that the decrease in the relative expression levels of t-SNARE proteins and probably other proteins may represent one of the putative mechanisms involved in this normalization and speculate that this also contributes to the change of glutamatergic synaptic response to the stimulation at 10 and 30Hz.

Thus, our data, as well as those of other laboratories, show how complex the functional consequences of Munc18.1 deficiency are, both biochemically and electrophysiologically, and to which a developmental parameter is now added. This shows how fundamental it is to restore the expression level/function of STXBP1 at a very early stage of development to prevent the triggering of all the cascade of events leading to long term dysfunction of cortical circuit. To this end, the use of chemical chaperones augmenting Munc18.1 level could represent promising treatment (Guiberson et al., 2018).

## Acknowledgments

We would also like to thank Francesca Bader for genotyping the mice (PBMC, platform headed by Emilie Pallesi-Pocachard at INMED) and technicians working at the INMED’s animal facility (plateform headed by Severine Pellegrino) for excellent technical support. Finally, we thank Dr Jerôme Epsztein for his constructive remarks on the manuscript

## Author contributors

L.P., H.B., M.B.H., N.M.B. performed electrophysiological recordings and contributed to data analysis. E.P.P and A.M. performed western blot experiments; M.M. and P.P.L.S contributed to the discussion of the data. L.A. designed the work, performed electrophysiological recordings, analyzed data and wrote the manuscript.

